# Cytotoxic CD4^+^ T cells eliminate senescent cells by targeting cytomegalovirus antigen

**DOI:** 10.1101/2023.03.06.531282

**Authors:** Tatsuya Hasegawa, Tomonori Oka, Heehwa G. Son, Valeria S. Oliver-García, Marjan Azin, Thomas M. Eisenhaure, David J. Lieb, Nir Hacohen, Shadmehr Demehri

**Affiliations:** Center for Cancer Immunology, Center for Cancer Research, Massachusetts General Hospital and Harvard Medical School, Boston, MA, USA; Cutaneous Biology Research Center, Department of Dermatology, Massachusetts General Hospital and Harvard Medical School, Boston, MA, USA; Shiseido Global Innovation Center, Yokohama, Japan; Broad Institute of MIT and Harvard, Cambridge, MA, USA

**Keywords:** Cytotoxic CD4^+^ T cell, senescent cell, cytomegalovirus, glycoprotein B, fibroblast, aging, skin

## Abstract

Senescent cell accumulation has been implicated in the pathogenesis of aging-associated diseases including cancer. The mechanism that prevents the accumulation of senescent cells in aging human organs is unclear. Here, we demonstrate that a virus-immune axis controls the senescent fibroblast accumulation in the human skin. Senescent fibroblasts increased in old compared with young skin. However, they did not increase with advancing age in the elderly. Increased CXCL9 and cytotoxic CD4^+^ T cell (CD4 CTL) recruitment were significantly associated with reduced senescent fibroblasts in the old skin. Senescent fibroblasts expressed human leukocyte antigen class II (HLA-II) and human cytomegalovirus glycoprotein B (HCMV-gB), becoming direct CD4 CTL targets. Skin-resident CD4 CTL eliminated HCMV-gB^+^ senescent fibroblasts in an HLA-II-dependent manner, and HCMV-gB activated CD4 CTL from the human skin. Collectively, our findings demonstrate HCMV reactivation in senescent cells, which can be directly eliminated by CD4 CTL through the recognition of the HCMV-gB antigen.

## INTRODUCTION

Senescent cells, which develop in response to cellular stress, exhibit irreversible arrest in proliferation while resisting death and can accumulate in the body with age (Di Micco et al., 2021; He and Sharpless, 2017). Despite their permanent cell cycle arrest, senescent cells are not inert. They actively communicate with their surroundings and influence the tissue microenvironment through multiple secretary molecules including pro-inflammatory cytokines and tissue-remodeling factors that are collectively termed the senescence-associated secretory phenotype (SASP) (Mahmoudi et al., 2019). Thus, senescent cells fuel a chronic inflammatory state in tissues, which in turn, leads to the development of cancer and aging-associated degenerative disorders (Kirkland and Tchkonia, 2017).

Accumulating evidence demonstrates that genetic or pharmacological approaches to eliminate senescent cells from aging tissues can restore tissue homeostasis and lead to increased healthy lifespan in mice (Mahmoudi et al., 2019; Paez-Ribes et al., 2019). However, the current genetic and pharmacological approaches produce substantial side effects and lack long-term durability (He and Sharpless, 2017; Kirkland and Tchkonia, 2017). Considering that the senescent cells produce SASP, their immunogenic phenotype marks them as potential targets for surveillance and clearance by the immune system. However, the immune clearance of senescent cells is hampered by the immunomodulatory molecules expressed by the senescent cells and the immunosuppressive factors in their microenvironment in the experimental mouse models (Kang et al., 2011; Kansara et al., 2013; Krizhanovsky et al., 2008; Pereira et al., 2019). More importantly, it remains unclear how immunity against senescent cells is regulated in humans.

The development and phenotype of senescent cells fundamentally differ between mice and humans including the role of telomere shortening and oxidative stress in the induction of cellular senescence (Coppe et al., 2010; Itahana et al., 2004; Parrinello et al., 2003; Wadhwa et al., 2002; Wright and Shay, 2000). A notable limitation of experimental mouse models to study senescence relates to their inability to fully capture the spectrum of immune responses against senescent cells due to their lack of pathogen exposure and evolutionarily distinct immunosurveillance mechanisms (Carr et al., 2016; Mestas and Hughes, 2004; Pulendran and Davis, 2020). Therefore, we explored the mechanism of senescent cell clearance in humans and discovered a novel commensal virome-immune axis that prevents the accumulation of senescent cells in aging skin.

## RESULTS

### Senescent cells are increased in the old human skin

We screened over 800 human skin samples from various ages and anatomical sites and identified cohorts of sun-protected truncal skin from young (n = 23, average age: 23.1) and old (n = 31, average age: 62.1) women (Supplementary Table 1). Reduced epidermal thickness in the old versus young skin demonstrated the biological evidence of skin aging in the selected skin samples (Supplementary Table 1). These skin cohorts enabled the evaluation of age-associated senescent cell accumulation in a human organ while excluding confounding factors like ultraviolet (UV) radiation, anatomic site variations, gender differences, and hair follicle density (Supplementary Table 1). To determine whether the number of senescent cells changed with age, we stained human skin samples with p16^INK4a^, a marker of cellular senescence (He and Sharpless, 2017; Ressler et al., 2006). The number of p16^INK4a^ positive cells was significantly increased in the epidermis and dermis with age; however, the magnitude of this increase was significantly higher in the dermis (Fig. 1A-C). Over 80% of dermal p16^INK4a^ positive cells expressed vimentin (Supplementary Fig. 1A), indicating that most of the senescent cells in the old human dermis were fibroblasts. Accordingly, the number and percentage of p16^INK4a+^ vimentin^+^ fibroblasts were significantly increased in the old compared with young skin (Fig. 1D, E and Supplementary Fig. 1B). Likewise, the number of dermal senescent cells was positively correlated (*r* = 0.5898) with age across the young and old skin samples (*P* < 0.0001; Fig. 1F). However, the number of dermal senescent cells did not show any significant correlation (*r* = 0.2121) with advancing age within the old skin cohort (*P* = 0.2520; Fig. 1G). This unexpected finding suggests that biological factor(s) other than increasing age may govern the accumulation of senescent cells in the elderly.

**Figure 1.**
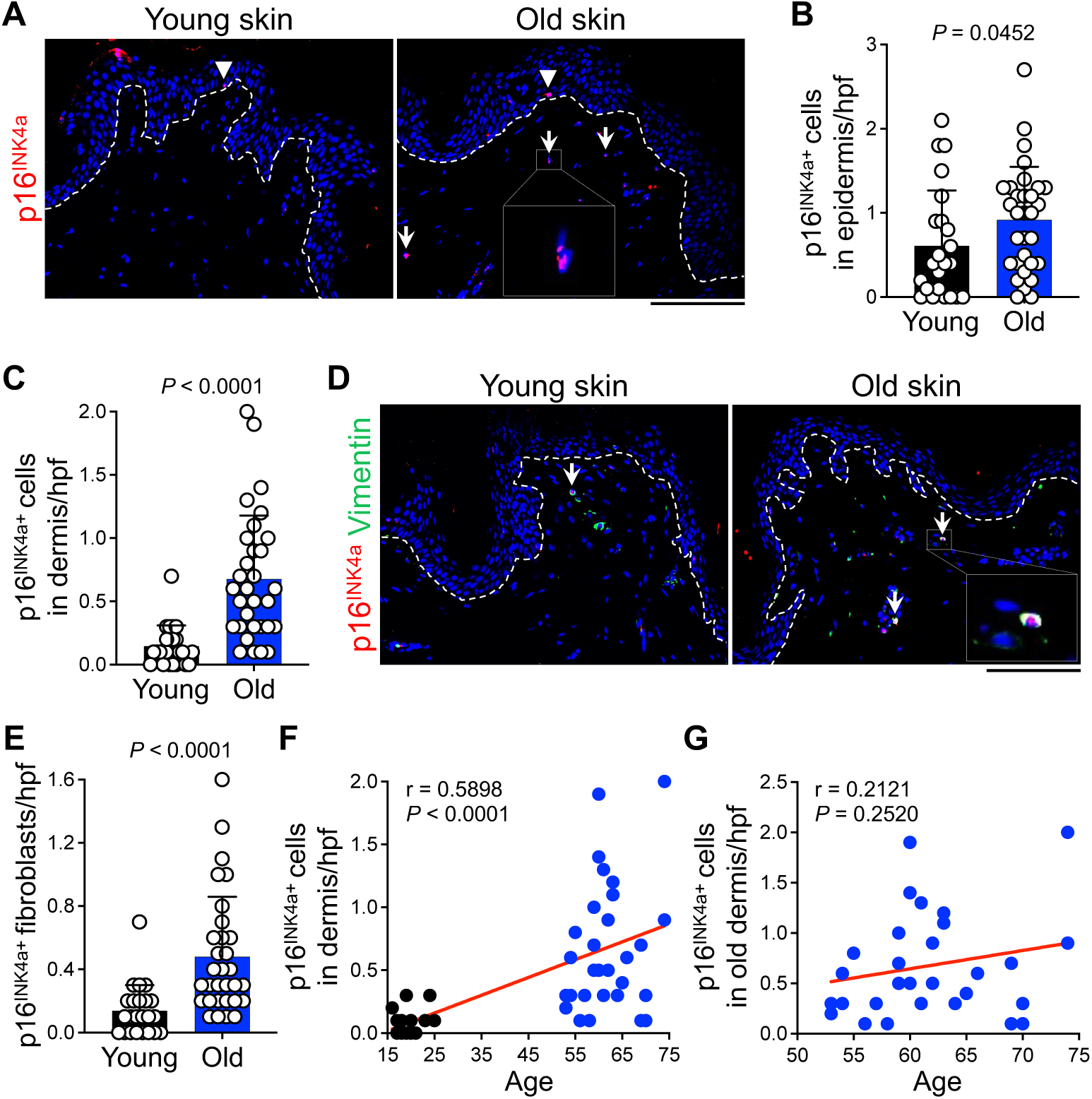
Senescent cell accumulation in the old human skin is not correlated with increasing age. (**A**) Representative immunofluorescence (IF) staining of p16^INK4a^ (red) in young and old human skin samples. Arrowheads point to p16^INK4a^ positive cells in the epidermis and arrows point to p16^INK4a^ positive cells in the dermis. (**B** and **C**) Quantification of p16^INK4a^ positive senescent cells in the epidermis (B) and dermis (C) per high power field (hpf) (Mann-Whitney *U* test). (**D**) Representative IF staining of p16^INK4a^ (red) and vimentin (green) in young and old skin samples. Arrows point to p16^INK4a+^ Vimentin^+^ fibroblasts in the dermis. (**E**) Quantification of p16^INK4a+^ Vimentin^+^ fibroblasts per hpf (Mann-Whitney *U* test). (**F**) Correlation between the number of dermal senescent cells and age across young and old skin samples (Student’s *t*-test for the Pearson correlation coefficient). (**G**) Correlation between the number of dermal senescent cells and age within the old skin samples (Student’s *t*-test for the Pearson correlation coefficient). Nuclei are stained with DAPI (blue). Dotted lines in IF images mark the epidermal basement membrane. Cells are counted blindly and averaged across 10 randomly selected hpf per skin sample. Bar graphs show mean + SD. *n* = 23 in young and *n* = 31 in the old skin group. Scale bars: 100 μm.

### CD4 CTL are the dominant cytotoxic lymphocytes in the aging human skin

To identify the factor(s) that regulate senescent cells accumulation in the old skin, we examined whether epidermal condition, blood or lymphatic vessel density impacted senescent cell numbers. Epidermal thickness, blood and lymphatic vessel density were significantly reduced in the old versus young skin samples; however, these factors had no significant correlation with the number of dermal senescent cells in the old skin samples (Supplementary Fig. 2). A comprehensive skin immune cell profiling on tissue sections revealed that several innate and adaptive immune cell types were highly enriched in the old dermis (Fig. 2A and Supplementary Fig. 3). Importantly, the number of several cytotoxic immune cell types was negatively correlated with the number of dermal senescent cells in the old skin samples (Supplementary Table 2). Among them, the number of cytotoxic CD4^+^ T cells (CD4 CTL), which were marked by perforin expression, was most negatively correlated (*r* = −0.6796) with the number of dermal senescent cells in the old skin samples (*P* < 0.0001; Fig. 2B). The number of perforin^+^ CD4 CTL was significantly increased in the old compared with young dermis (Fig. 2C, D). Notably, 83.7% of perforin^+^ dermal cells in the old skin samples were CD4^+^ T cells (Fig. 2E), indicating that CD4 CTL were the dominant cytotoxic lymphocytes responsible for immunosurveillance in the old human skin. These findings suggest that CD4 CTL are the effector cells responsible for the clearance of senescent cells in the old human skin.

**Figure 2.**
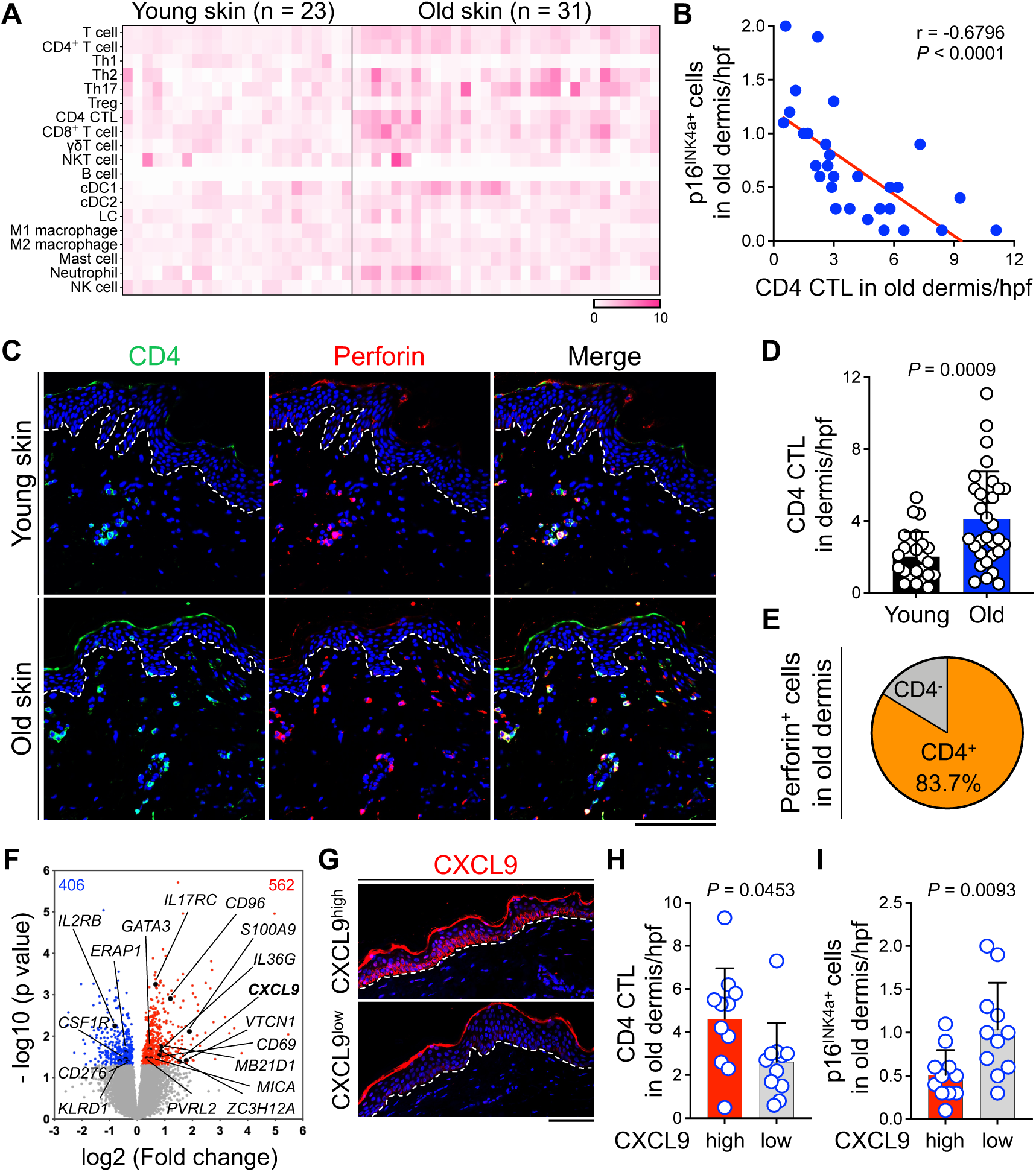
CD4 CTL are the dominant cytotoxic lymphocytes in the human skin, and their number is inversely correlated with the number of dermal senescent cells in the old skin. (**A**) Heatmap of aging-associated changes in the number of dermal immune cells, determined by multiplex immunostaining on tissue sections. The color intensity represents a relative change compared with the young group. *n* = 23 in young and *n* = 31 in the old skin group. (**B**) Correlation between the number of dermal senescent cells and dermal CD4 CTL in old skin samples (*n* = 31 old skin samples, Student’s *t*-test for the Pearson correlation coefficient). (**C**) Representative IF staining of CD4 (green) and Perforin (red) in young and old human skin samples. Note that CD4^+^ cells are CD3^+^ T cells. (**D**) Quantification of dermal CD4^+^ Perforin^+^ CD4 CTL per hpf (*n* = 23 in young and *n* = 31 in old skin group, Mann-Whitney *U* test). (**E**) Pie chart showing the percentage of Perforin^+^ cells in the old dermis (*n* = 31 old skin samples). (**F**) Volcano plot of RNA-Seq data displaying the gene expression pattern in the old versus young human skin samples. Significantly upregulated genes are highlighted in red, and downregulated genes are highlighted in blue (*P* < 0.05 is considered significant). Selected immune-related genes based on cluster analysis are indicated. (**G**) Representative IF staining of CXCL9 (red) in the old human skin samples. (**H**) Quantification of dermal CD4 CTL per hpf in CXCL9^high^ versus CXCL9^low^ group of old skin samples (*n* = 11 in each group, Mann-Whitney *U* test). (**I**) Quantification of p16^INK4a^ positive dermal senescent cells per hpf in CXCL9^high^ versus CXCL9^low^ group of old skin samples. (*n* = 11 in each group, Mann-Whitney *U* test). Nuclei are stained with DAPI (blue). Dotted lines in IF images mark the epidermal basement membrane. Cells are counted blindly and averaged across 10 randomly selected hpf per skin sample. Bar graphs show mean + SD. Scale bars: 100 μm.

To determine the factors that recruited CD4 CTL in the old skin, we performed RNA-sequencing (RNA-Seq) of young and old skin samples. Cluster analysis showed that immune-related genes, including *CD69*, *CD96*, *CD276*, *CXCL9*, *KLRD1*, *IL2RB*, *IL17RC*, *IL36G*, *MB21D1*, *S100A9* and *VTCN1*, were among the genes significantly altered in the old versus young skin (Fig. 2F). Among these, we focused on CXCL9 chemokine, which is a ligand of CXCR3 expressed on CD4 CTL (Takeuchi et al., 2016). CXCL9 expressing cells were mainly localized in the epidermal basal layer of the old skin samples (Fig. 2G). Dividing the old skin samples into CXCL9^high^ and CXCL9^low^ groups revealed that CXCL9^high^ skin contained a significantly higher number of CD4 CTL and a lower number of dermal senescent cells compared with CXCL9^low^ skin (Fig. 2H, I). These findings indicate that CXCL9 expressed by keratinocytes recruits CD4 CTL to the skin, which may result in the clearance of dermal senescent cells.

### Senescent fibroblasts are directly targeted by CD4 CTL

To examine whether CD4 CTL can target senescent fibroblasts, we generated senescent human dermal fibroblasts *ex vivo* through repeated passage-induced replicative senescence, which mimics cellular senescence in the sun-protected old skin (Campisi, 1998). Senescence-associated β-galactosidase (SA-β-Gal) staining marked the senescent fibroblasts (Fig. 3A). RNA-Seq analysis further confirmed the cellular senescence gene set enrichment and revealed that several ligands recognized by cytotoxic lymphocytes were significantly altered in replication-induced senescent compared with normal fibroblasts (Fig. 3B and Supplementary Fig. 4A). Among these, UL16 Binding Protein 2 (ULBP2), which is a ligand for activating natural killer group 2D (NKG2D) receptor on cytotoxic lymphocytes (Gonzalez et al., 2008), was highly upregulated in senescent fibroblasts (Fig. 3B). Accordingly, ULBP2 was expressed on the surface of senescent fibroblasts in contrast to its lack of expression on normal fibroblasts (Fig. 3C, D). We detected the expression of ULBP2 in the senescent fibroblasts in old human skin samples (Supplementary Fig. 4B). Human leukocyte antigen class II (HLA-II) surface expression is pivotal for CD4 CTL-elicited immunity. To determine whether senescent fibroblasts could be direct CD4 CTL targets, we examined HLA-II expression on senescent fibroblasts. Importantly, we found that HLA-II was highly expressed on senescent fibroblasts while its expression was negligible on normal fibroblasts (Fig. 3E, F). Consistent with this finding, the number of HLA-II^+^ senescent fibroblasts was significantly increased in the old compared with young human skin (Fig. 3G, H). UVA radiation is a prominent cause of skin aging (Lan et al., 2019), and is known to induce senescence in fibroblasts (Lan et al., 2019). We used UVA radiation to generate senescent human dermal fibroblasts *ex vivo* (Supplementary Fig. 4C). ULBP2 and HLA-II levels were markedly increased on the surface of UVA-induced senescent fibroblasts (Supplementary Fig. 4D-K). These findings demonstrate that senescent fibroblasts are highly immunogenic targets, which can be directly recognized by CD4 CTL in the human skin.

**Figure 3.**
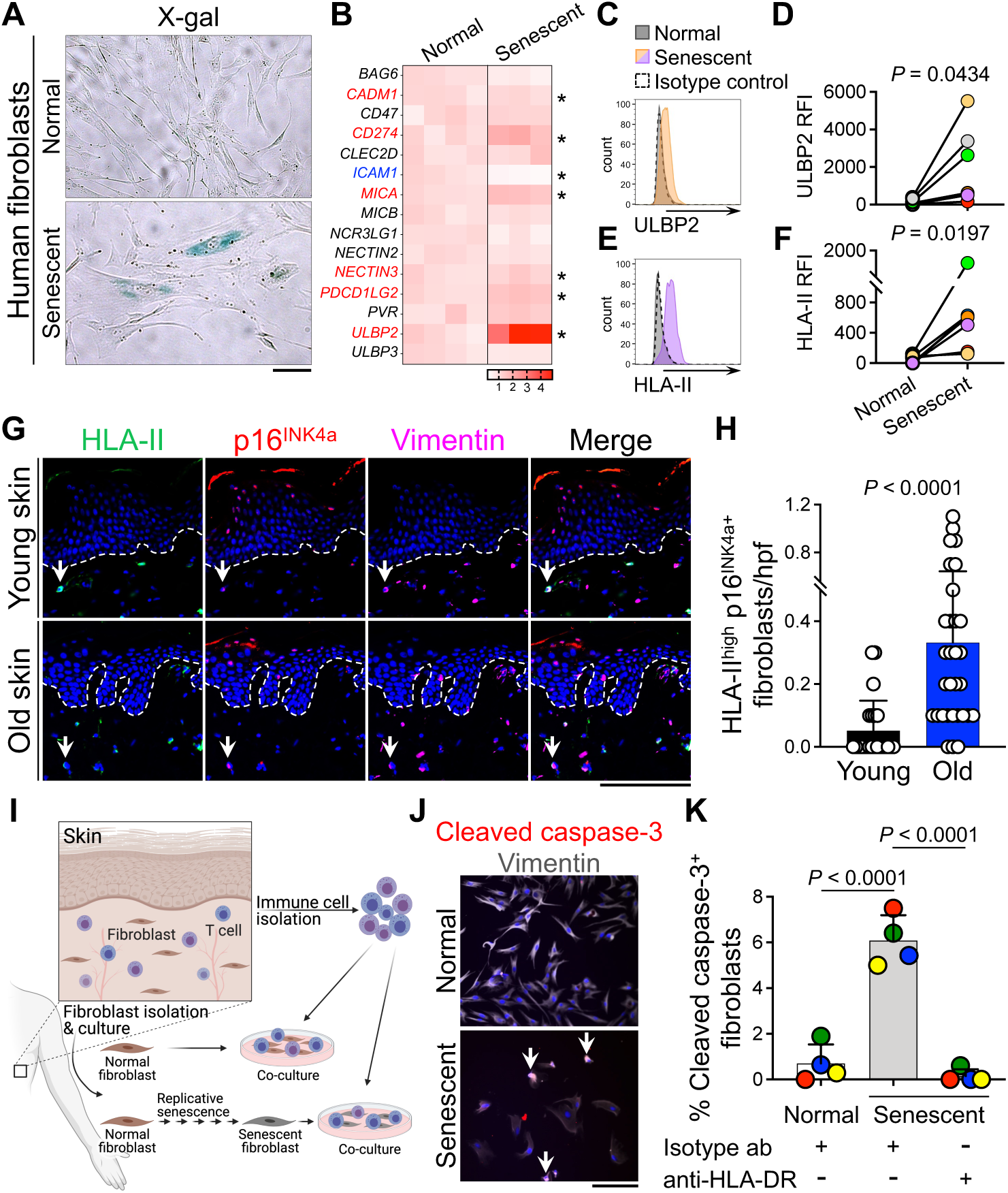
Skin-resident T cells eliminate senescent fibroblasts in an HLA-II-dependent manner. (**A**) Representative senescence-associated β-galactosidase (SA-β-Gal) staining of normal and senescent dermal fibroblasts. (**B**) Heatmap representing expressions of immune cell-activating ligands genes from RNA-Seq data. Blue indicates downregulation (log_2_ Fold change < −0.4, \**P* < 0.05), and red indicates upregulation (log_2_ Fold change > 0.4, \**P* < 0.05) in senescent (*n* = 3) versus normal fibroblasts (*n* = 4). (**C** and **D**) Representative flow cytometry histogram (C) and relative fluorescence intensity (RFI) (D) of ULBP2 on the surface of normal and senescent fibroblasts (*n* = 7 pairs of old human dermal fibroblasts (average age: 53.7), paired *t*-test). (**E** and **F**) Representative flow cytometry histogram (E) and RFI (F) of HLA-II on the surface of normal and senescent fibroblasts (*n* = 7 pairs of old human dermal fibroblasts (average age: 53.7), paired *t*-test). (**G**) Representative IF staining of HLA-II (green), p16^INK4a^ (red), and vimentin (magenta) in the young and old skin samples. Arrows point to HLA-II^high^ senescent fibroblasts in the dermis. Dotted lines mark the epidermal basement membrane. (**H**) Quantification of HLA-II^high^ senescent fibroblasts in the skin per hpf (*n* = 23 in young and *n* = 31 in old skin group, Mann-Whitney *U* test). (**I**) Experimental scheme to assay autologous immune cell-elicited cytotoxicity against senescent fibroblasts. Schematic diagram was created with biorender.com. (**J**) Representative immunocytochemistry (ICC) staining of cleaved caspase-3 (red) and vimentin (gray) in normal and senescent fibroblasts after co-culture with skin-derived autologous immune cells. Arrows point to cleaved caspase-3^+^ apoptotic fibroblasts. (**K**) Frequency of cleaved caspase-3^+^ normal and senescent fibroblasts co-cultured with skin-derived autologous immune cells. Fibroblasts were pre-incubated with pan-HLA-DR blocking or isotype control antibodies. Skin immune cell to fibroblast ratio of 50:1 (*n* = 4 individual old donors, one-way ANOVA). Nuclei are stained with DAPI (blue). Cells are counted blindly and averaged across 10 randomly selected hpf per sample. Bar graphs show mean + SD. Scale bars: 100 μm.

### CD4 CTL eliminate senescent fibroblasts in an HLA-II-dependent manner

To determine whether CD4 CTL can kill senescent fibroblasts, we co-cultured normal and replication-induced senescent fibroblasts with immune cells isolated from the old human skin. The skin-derived allogeneic T cells caused apoptosis specifically in the senescent fibroblasts in an HLA-II-dependent manner (Supplementary Fig. 5). Next, we established a co-culture system in which normal and senescent fibroblasts were exposed to autologous T cells that were isolated from the same human skin (Fig. 3I). Autologous T cells induced cleaved caspase-3^+^ apoptotic senescent fibroblasts at a significantly higher ratio compared with normal fibroblasts (Fig. 3J, K). Importantly, HLA-DR antibody blockade abolished the skin T cell-induced apoptosis of senescent fibroblasts (Fig. 3K). These findings demonstrate that senescent fibroblasts are eliminated by T cells in an HLA-II-dependent manner.

### CD4 CTL eliminate senescent fibroblasts by targeting HCMV-gB antigen

Next, we investigated which HLA-II-bound antigen triggered CD4 CTL killing of senescent fibroblasts. Commensal human cytomegalovirus (HCMV) derived-glycoprotein B (gB) is a principal target of CD4^+^ T cells and can activate CD4 CTL to eliminate host cells in humans (Hegde et al., 2005). Furthermore, gB-specific CD4^+^ T cells are found in the blood of 95% of healthy individuals (Pachnio et al., 2015). HCMV DNA and RNA were detectable in the normal human skin, and their levels were significantly increased in the old compared with young skin samples (Supplementary Fig. 6A-D). HCMV RNA expression was particularly evident in the dermal fibroblasts in the old skin (Supplementary Fig. 6D, E). Strikingly, we found that senescent fibroblasts expressed HCMV-gB in the human skin and the number of HCMV-gB^+^ senescent fibroblasts was significantly increased in the old compared with young skin (Fig. 4A, B). In line with the findings *in vivo*, replication- and UVA-induced senescent fibroblasts highly upregulated HCMV DNA and HCMV-gB expression *ex vivo* (Fig. 4C, D and Supplementary Fig. 7A-C). HCMV induces the accumulation of early endosomes in the infected cells (Karleusa et al., 2018; Lucin et al., 2020), and HCMV-gB is sorted into endosomes and presented on HLA-II (Hegde et al., 2005). Similar to HCMV-infected fetal fibroblasts, we found that HCMV-gB trafficked into endosomes of replication- and UVA-induced adult senescent fibroblasts (Supplementary Fig. 7D-F). These findings reveal that HCMV is activated in senescent fibroblasts and HCMV-gB can be displayed as an endogenous antigen on HLA-II.

**Figure 4.**
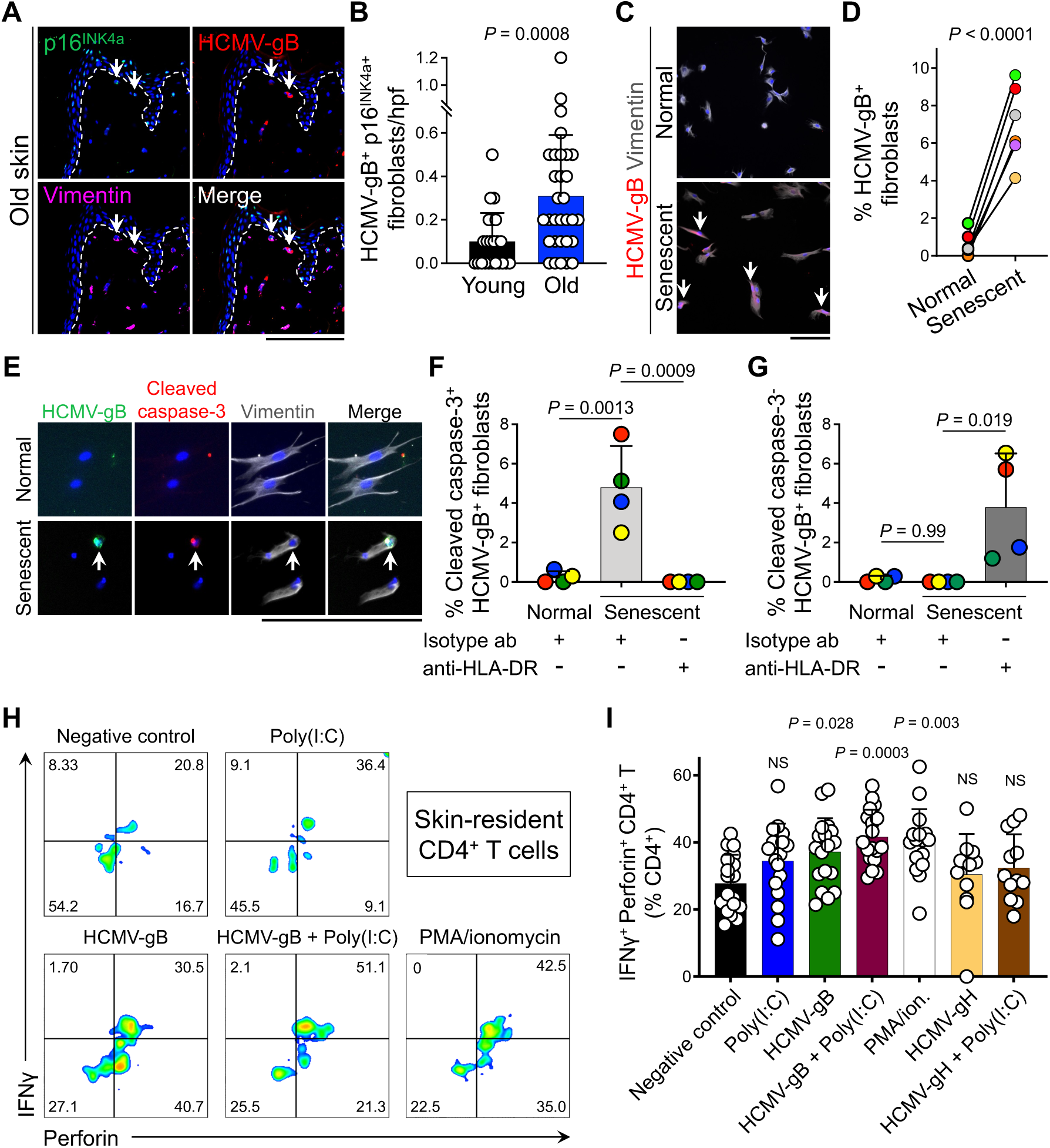
HCMV-gB antigen is expressed by senescent fibroblasts and activates skin-resident CD4 CTL. (**A**) Representative IF staining of p16^INK4a^ (green), HCMV-gB (red), and vimentin (magenta) in the old human skin. Arrows point to HCMV-gB^+^ senescent fibroblasts. Dotted lines mark the epidermal basement membrane. (**B**) Quantification of HCMV-gB^+^ p16^INK4a+^ fibroblasts in dermis per hpf (*n* = 23 in young and *n* = 31 in old skin group, Mann-Whitney *U* test). (**C**) Representative ICC staining of HCMV-gB (red) and vimentin (gray) in normal and senescent fibroblasts. Arrows point to HCMV-gB^+^ fibroblasts. (**D**) Frequency of HCMV-gB^+^ normal or senescent fibroblasts (*n* = 7 pairs of adult human dermal fibroblasts, paired *t*-test). (**E**) Representative ICC staining of HCMV-gB (green), cleaved caspase-3 (red), and vimentin (gray) in normal and senescent fibroblasts after co-culture with skin-derived autologous immune cells. Arrows point to an HCMV-gB^+^ apoptotic fibroblast. (**F** and **G**) Frequency of cleaved caspase-3^+^ HCMV-gB^+^ (F), cleaved caspase-3^-^ HCMV-gB^+^ (G) normal and senescent fibroblasts co-cultured with skin-derived autologous immune cells. Fibroblasts were pre-incubated with pan-HLA-DR blocking or isotype control antibodies. Skin immune cell to fibroblast ratio of 50:1 (*n* = 4 individual donors, one-way ANOVA). (**H** and **I**) Representative flow cytometry plots (H) and quantification (I) of skin-resident IFNγ^+^ Perforin^+^ CD4^+^ T cells following stimulation of recombinant HCMV-gB, HCMV-gH and/or poly(I:C). The percentage of CD4^+^ T cells in each quadrant is shown on the flow cytometry plots. Stimulation with phorbol-12-myristate-13-acetate and ionomycin (PMA/ion.) was used as a positive control (*n* = 18 in negative control, *n* = 18 in Poly(I:C), *n* = 18 in HCMV-gB, *n* = 18 in HCMV-gB + Poly(I:C), *n* = 16 in PMA/Ionomycin, *n* = 12 in HCMV-gH, and *n* = 12 in HCMV-gH + Poly(I:C) group, one-way ANOVA, Dunnett’s multiple comparison test). Skin samples from five independent donors were used to isolate skin-resident immune cells. Nuclei are stained with DAPI (blue). Cells are counted blindly and averaged across 10 randomly selected hpf per sample. Bar graphs show mean + SD. Scale bars: 100 μm.

Autologous T cell/fibroblast co-culture assay revealed that most of the apoptotic senescent fibroblasts expressed HCMV-gB (Fig. 4E, F). Importantly, HCMV-gB^+^ senescent fibroblasts remained intact upon HLA-DR blockade (Fig. 4G). Finally, recombinant HCMV-gB plus Poly(I:C) induced interferon-γ (IFNγ) expression in perforin^+^ CD4 CTL from the human skin *ex vivo* while HMCV glycoprotein H (gH) plus Poly(I:C) did not induce IFNγ expression in CD4 CTL (Fig. 4H, I). Together, these findings indicate that HCMV-gB-specific CD4 CTL contribute to the clearance of senescent fibroblasts in the aging human skin (Supplementary Fig. 8).

## DISCUSSION

Our findings reveal a previously unknown role for CD4 CTL in the clearance of senescent cells that experience commensal cytomegalovirus reactivation. These findings have significant implications for understanding how the immunity against senescent cells is regulated in humans. Senescent cells, mainly dermal fibroblasts, are accumulated in the old compared with young skin; however, their number does not linearly correlate with advancing age in elderly. Among the possible explanations for this age-independent accumulation of senescent cells, we conclude that CD4 CTL is a key regulator of senescent cells in the old skin. Cytotoxic lymphocytes utilize the perforin/granzyme pathway to kill virally infected and tumor cells (Trapani and Smyth, 2002). Unexpectedly, we find that CD4^+^ T cells are the largest population of perforin-expressing cytotoxic lymphocytes in the old skin, and their frequency in the dermis most negatively correlates with the accumulation of senescent cells in old human skin. Examination of the immunological nature of senescent fibroblasts as targets of CD4 CTL reveal that replication and UVA-induced senescent human fibroblasts upregulate HLA-II and HCMV-gB expression. Endogenous HCMV-derived gB is sorted to endosomes and presented on HLA-II to CD4^+^ T cells (Hegde et al., 2005). CD4 CTL from the human skin specifically eliminate HCMV-gB^+^ senescent fibroblasts in an HLA-II-dependent manner.

CD4 CTL are detected mostly during viral infection in humans, where their direct anti-viral effector functions to help control the infection (Oh and Fong, 2021; Takeuchi and Saito, 2017). Notably, the high presence of circulating CD4 CTL is a prominent characteristic of supercentenarians, who live more than 110 years in good health due to the delayed onset of age-related diseases and reduced morbidity (Hashimoto et al., 2019). It is intriguing to postulate that, in addition to fighting infection, CD4 CTL contribute to the elimination of senescent cells and their associated diseases in supercentenarians to achieve exceptional longevity.

Although HCMV is an infectious cause of birth defects and can cause serious morbidity in severely immunocompromised individuals (Griffiths and Reeves, 2021; Rawlinson et al., 2017), it is a pervasive herpesvirus that establishes lifelong latent infection in the majority of the human population without any symptoms in healthy hosts (Shenk and Alwine, 2014). HCMV produces several immunodominant antigens and has a profound influence on shaping the repertoire of the adaptive immunity in healthy individuals during aging (Brodin et al., 2015). HCMV-specific CD4 CTL display high cytotoxicity while producing a low amount of cytokines, which enables them to effectively fight HCMV reactivation whilst minimizing tissue inflammation (Parry et al., 2021). We find that HCMV is reactivated upon the induction of cellular senescence. This key discovery explains how CD4 CTL, and likely CD8 CTL and NK cells, can detect the senescent fibroblasts in the mix of healthy cells, which are also infected with HCMV as a commensal virus. Our finding is supported by the observation that HCMV upregulates p16^INK4a^ to increase its replication (Noris et al., 2002). HCMV reactivation upon replication- and UVA-induced senescence, coupled with HLA-II and stress ligands induction on the senescent cells, enables the HCMV-specific CD4 CTL to effectively target and eliminate the senescent cells.

Our findings provide a novel insight into the complex human immune system-commensal virome interactome. Our research indicates that the human immune system has evolved to establish a symbiotic relationship with HCMV that prevents the accumulation of senescent cells during aging. Thus, HCMV can have a beneficial impact on human health, which is mediated by a competent anti-viral T cell immunity. We propose a novel strategy for senescent cell clearance through vaccination against commensal HCMV antigens in order to boost anti-HCMV T cell immunity and prevent aging and aging-associated diseases.

## METHODS

### Human skin tissue study

The study of deidentified normal adult human skin samples was reviewed and approved by the Institutional Review Board of Massachusetts General Hospital. Patient consent for experiments was not required because US laws consider human tissue left over from surgery as discarded material.

### Isolation and culture of human dermal fibroblasts

Dermal fibroblasts were isolated from discarded normal skin samples, which were generated as part of the surgery. Subcutaneous fat tissue was removed from human skin tissue, and the tissue pieces were incubated in dispase solution (Stemcell, Vancouver, Canada, 07913) overnight at 4°C. After digestion, the epidermis was separated from the dermis. The obtained dermis was incubated in collagenase/hyaluronidase (Stemcell, 07912) overnight at 37°C. Fibroblasts were collected through a 70 µm cell strainer and were seeded at a density of 3-5 x 10^4^ cells/cm^2^ into 75 cm^2^ cell culture flasks, and cultured in DMEM medium (Thermo Fisher Scientific, Waltham, MA 11-965-118), including 10% fetal bovine serum (FBS), 1% penicillin/streptomycin, and 1% glutamine, at 37°C under an atmosphere of 5% CO_2_ in the air. Human fetal dermal fibroblasts (ScienCell Research Laboratories, Carlsbad, CA, 2300) and neonatal dermal fibroblasts (Lonza, Basal, Switzerland, CC-2509) were purchased. Cells were cultured as described above.

### Human skin immune cell isolation

Immune cells were isolated from human skin as previously described (Watanabe et al., 2015). Briefly, discarded normal skin samples generated as part of surgery were obtained. Subcutaneous fat tissue was removed from human skin tissue, and the remaining tissue was minced. Skin tissues were minced and digested in RPMI 1640 medium (Thermo Fisher Scientific, 21-870-092) including 0.05% DNase-I (Sigma-Aldrich, St. Louis, MO, 10104159001) and 0.2% collagenase-I (Thermo Fisher Scientific, LS004196) for 2 h at 37°C. Then cells were collected through a 70 µm cell strainer and were incubated in RPMI 1640 medium including 20% FBS, 1% penicillin/streptomycin, 1% glutamine, 0.00035% β-mercaptoethanol, and 2 ng/ml human interleukin (IL)-2 recombinant (BioLegend, San Diego, CA, 589104).

### Histology

Human skin samples were fixed with 4% paraformaldehyde (PFA) and embedded in paraffin. 5 µm sections were cut and deparaffinized. After being permeabilized with 0.2% Triton-X (Thermo Fisher Scientific, BP151) in phosphate-buffered saline (PBS), antigen retrieval was performed using a pressure cooker in antigen unmasking solution (Vector Laboratories, Burlingame, CA, H-3300-250) for 20 min. Slides were washed three times for 5 min each in PBS including 0.1% Tween 20 (Sigma-Aldrich, P1379). Slides were blocked with 5% normal goat serum (Sigma-Aldrich, G9023) and 5% bovine serum albumin (Thermo Fisher Scientific, BP1600) in PBS for 1 h. Slides were stained overnight at 4°C with primary antibodies (Supplementary Table 3A) diluted in the blocking buffer. Following primary antibody application, slides were washed and incubated in secondary antibodies (Supplementary Table 3A) diluted in the blocking buffer for 2 h at room temperature. Slides were washed and stained with 4’,6-diamidino-2-phenylindole (DAPI, Invitrogen, D3571, 1:4000) in PBS for 10 min at room temperature. Slides were washed and mounted with ProLong Gold Antifade Reagent (Thermo Fisher Scientific, P36930). The stained tissues were imaged with a ZEISS confocal microscope (Zeiss, Oberkochen, Germany). Manual counting was performed using the ZEN Blue Software (Zeiss). Cell counts were reported as the average number of cells across 10 randomly selected high power fields (hpf, 200x magnification) per skin sample in each group. For hematoxylin and eosin staining, slides were stained according to standard procedures and mounted with Cytoseal XYL (Thermo Fisher Scientific, 8312-4). Whole-slide imaging was performed using a Zeiss Axio Scan.Z1 (Zeiss).

### Immunocytochemistry

Cells were cultured on chamber slide glasses (CELLTREAT Scientific Products, Pepperell, MA, 229168) and fixed with 4% PFA in 10 min at room temperature and were permeabilized with 0.2% Triton-X in PBS in 10 min at room temperature. Slides were washed three times for 5 min each in PBS. Slides were blocked with 5% normal goat serum and 5% bovine serum albumin in PBS for 30 min. Slides were stained overnight at 4°C with primary antibodies (Supplementary Table 3A) diluted in the blocking buffer. Following primary antibody application, slides were washed and incubated in secondary antibodies (Supplementary Table 3A) diluted in the blocking buffer for 1 h at room temperature. Slides were washed as above and stained with DAPI in PBS for 10 min at room temperature. Slides were washed as above and mounted with ProLong Gold Antifade Reagent. Stained cells were imaged with a ZEISS confocal microscope. Cell counts were reported as the average number of cells across 10 randomly selected hpf per well in each group.

### RNA-Seq analysis

Human skin tissues were homogenized with RLT buffer (Qiagen, Hilden, Germany, 79216) supplemented with 1% β-mercaptoethanol (Thermo Fisher Scientific, 21-985-023). Full-length cDNA and sequencing libraries were prepared from 1 ng RNA using the Smart-Seq2 protocol as previously described (Picelli et al., 2014). Libraries were sequenced on a Novaseq 6000 (Illumina) through the Broad Genomics Platform. The FASTQ files were aligned to the human genome/hg19 (GENCODE v19) by STAR-2.5.1b (Dobin et al., 2013). Aligned transcripts were quantified by using RSEM-1.2.3.1 (Li and Dewey, 2011). Differentially expressed genes (DEG) were analyzed by DESEq2(Love et al., 2014). Cultured fibroblasts were prepared in TCL buffer (Qiagen, 1031576) supplemented with 1% β-mercaptoethanol. Then, each sample was added into a 96-well Eppendorf twin-tec barcoded plate provided by the Broad Institute (Cambridge, MA). Modified SmartSeq2 complementary DNA and Illumina Nextera XT library construction and sequencing were conducted at the Broad Institute using the Illumina NextSeq 500 System. The quality of FASTQ files was examined using FastQC-0.11.8. The sequences were mapped to the human genome/GRCh38 using STAR-2.5.3 (Dobin et al., 2013). Sequences located at transcripts were quantified by using RSEM-1.3.1 (Li and Dewey, 2011). DEGs were analyzed using DESEq2-1.24.0 (Love et al., 2014). Original data are available in the NCBI Gene Expression Omnibus (GEO) with accession number GSE191055.

### Senescence-associated β-galactosidase (SA-β-Gal) staining

Cells were stained with β-galactosidase at pH 6.0 with Senescence β-galactosidase Staining Kit (Cell Signaling Technology, 9860), according to the manufacturer’s protocol. Stained cells were imaged with a ZEISS confocal microscope.

### Flow cytometry

Cells were washed once with PBS, including 5% newborn calf serum (Thermo Fisher Scientific, 26010074) and 0.01% sodium azide (Sigma-Aldrich, S2002-100G), and stained with antibodies (Supplementary Table 3B) on ice for 30 min followed by secondary antibody if needed (Supplementary Table 3). For intracellular staining, following with surface marker staining, cells were fixed and permeabilized using True-Nuclear Transcription Factor Buffer Set (BioLegend, 424401). Permeabilized cells were stained with antibodies (Supplementary Table 3) overnight at 4°C. Cells were washed and then examined by BD LSRFortessa X-20 flow cytometer (BD Bioscience, Billerica, MA). Data were analyzed using FlowJo software (BD Life Sciences, Franklin Lakes, NJ).

### Cytotoxicity assay on senescent fibroblasts

Normal and senescent fibroblasts were seeded at a density of 5×10^3^ cells/well into 24 well plates, and were cultured in DMEM, including 10% FBS, 1% penicillin/streptomycin, 1% glutamine, overnight at 37°C. Fibroblasts were pre-treated with 100 µg/ml HLA-II blocking antibody (Bio X Cell, Lebanon, NH, BE0306) or 100 µg/ml isotype IgG antibody in the absence of FBS overnight at 37°C. Skin immune cells were subsequently added to each fibroblasts-placed well and co-cultured in the presence of 20 ng/ml human IL-2 recombinant and 20 ng/ml human IL-15 recombinant (BioLegend, 570304) in 24 well plates (Ratio of 50:1 immune cell-to-fibroblast). Following 6 h of co-culture, remaining adherent fibroblasts were fixed and stained as described in immunocytochemistry.

### Recombinant virus protein stimulation on human skin T cells

Isolated human skin immune cells were treated with 10 µg/ml of recombinant HCMV-gB (Abcam, Cambridge, UK, ab43040) or 10 µg/ml of recombinant HCMV-gH (MyBioSource, San Diego, CA, MBS1138239) in the presence or absence of 1.67 µg/ml of Poly (I.C) (Thermo Fisher Scientific, tlrl-pic). After 20 h of incubation, Brefeldin A was added at the concentration of 5 µg/ml and cells were incubated for 4 h and collected for flow cytometric analysis. As a positive control, cells were treated with 81 nM phorbol 12-myristate 13-acetate (PMA) plus 1.34 µM Ionomycin (BioLegend, 423301) for 30 min. Cells were stained as described in flow cytometry.

### RNA in situ hybridization

RNA *in situ* hybridization was performed as previously described (Schiferle et al., 2021). Briefly, RNA *in situ* hybridization was performed on PFA-fixed paraffin-embedded tissue sections using the RNAscope 2.5 HD detection reagent protocol (Advanced Cell Diagnostics, Newark, CA) with accommodation to simultaneously stain for vimentin protein. 5 µm sections were baked at 60°C for 60 min. Slides were treated with xylene, followed by 100% ethanol, and allowed to dry. Slides were treated with hydrogen peroxide at room temperature for 10 min and then washed with deionized water. Antigen retrieval was performed with RNAscope Target Retrieval Reagent (Advanced Cell Diagnostics, 322000) using a pressure cooker for 15 min. Slides were incubated with DNase-I (Sigma-Aldrich, D5319-500UG) at 37°C for 30 min in a HybEZ Oven II (Advanced Cell Diagnostics, 321720) and then washed. RNAscope Protease Plus (Advanced Cell Diagnostics, 322331) treatment was applied at 40°C for 15 min. After target probe amplification and hybridization steps, sections were stained with Fast RED reagent (RNAscope 2.5 HD Detection Reagents-RED, Advanced Cell Diagnostics, 322360). For hematoxylin staining, slides were washed with deionized water and then stained with hematoxylin (Sigma-Aldrich, GHS132-1L) for 1 min, followed by staining with 0.02% ammonium hydroxide (Ricca Chemical Company, Arlington, TX, 642-16). For immunofluorescent staining, slides were washed with deionized water and then PBS including 0.1% Tween 20. Slides were blocked with 5% goat serum and 5% bovine serum albumin in PBS, including 0.1% Tween 20, for 1 h at room temperature. Slides were stained as described in histology.

### Quantification of HCMV DNA

Quantification of HCMV DNA in human skin samples and fibroblasts was performed by quantitative real-time PCR as described previously (Czech-Kowalska et al., 2021; Kasztelewicz et al., 2017). For fibroblasts, DNA was isolated using Quick-DNA/RNA Microprep Plus kit (Zymo Research, Irvine, CA, D7005) following the kit instructions. For human skin, DNA was isolated using a Direct-zol DNA/RNA Miniprep Kit (Zymo Research, R2080) following the kit instructions. Quantitative real-time PCR used the SYBRGreen format and HCMV primers detected the lower matrix phosphoprotein (*UL83*) gene. *GAPDH* was used as the internal control gene. Primer sets are described in Supplementary Table 3C. PCR was performed on the 7500 Real-Time PCR System (Applied Biosystems, Inc., Foster City, CA) in a total volume of 25 μL in the presence of 5 μL of DNA sample, 12.5 μL of SYBRGreen PCR MasterMix (Bio-Rad, Hercules, CA, 1725121) and 250 nM of each of the primers. The temperature profile was 95 °C for 10 min, 40 cycles at 95 °C for 15 s and 60 °C for 60 s. At the end of each run, a melting curve analysis was performed. The melting temperature range for HCMV DNA positive samples was 81.5 ± 0.5 °C. Relative expression of *UL83* was calculated by the *ΔΔ*Ct method.

### UVA irradiation

Fibroblasts were irradiated through PBS with UVA (5 or 10 J) generated by a UVP XX-Series Bench Lamp, 115V (Thermo Fisher Scientific, UVP95004208), and cultured for one day or five days. Sham irradiation was used as a negative control. Radiation intensity was measured using a light meter (InternationalLight Technologies, Peabody, MA, ILT2400).

### Human cytomegalovirus infection

Human cytomegalovirus, AD-169, was purchased from American Type Culture Collection (Manassas, VA, VR-538). Viral concentration was determined by plaque assay. Human dermal fetal fibroblasts were seeded at a density of 2-3 x 10^4^ cells/cm^2^ and cultured as described in the culture of human dermal fibroblasts. One day after seeding, culture media was replaced with DMEM with 0.1% bovine serum albumin, and the cells were infected with a multiplicity of infection (MOI) of 1. Five days after infection, DNA was isolated from the cells. Three days after infection, infected cells were stained for immunocytochemistry as described above.

### Statistical analysis

All bar graphs and dot plots show mean + SD. Two-tailed Mann-Whitney *U* test was used as the test of significance for cell counts in human skin histological analysis. A two-tailed paired *t*-test was used for the cultured dermal fibroblasts assays. The Student’s *t*-test for the Pearson correlation coefficient was used as the test of significance for the linear regression in the scatter plots. One-way ANOVA with Tukey’s multiple comparison test was used for the co-culture cytotoxic assay. One-way ANOVA with Dunnett’s multiple comparison test was used for the HCMV-gB stimulation assay. *P* value < 0.05 was considered significant.

## ACKNOWLEDGMENTS

Shadmehr Demehri, M.D., Ph.D., holds a Career Award for Medical Scientists award from the Burroughs Wellcome Fund. TH was supported by Shiseido Co. Ltd. The authors gratefully acknowledge support from Shiseido Co. Ltd.

## AUTHOR CONTRIBUTIONS

TH and SD conceived the study. TH, TO and SD designed the experiments. TH, TO, HGS, VSO and MA performed the experiments and analyzed the data. TME, DJL, and NH contributed experimental data. TH, TO and SD interpreted the data. TH, TO and SD wrote the manuscript.

## Supplementary Materials

### SUPPLEMENTARY FIGURES

**Supplementary Figure 1.**
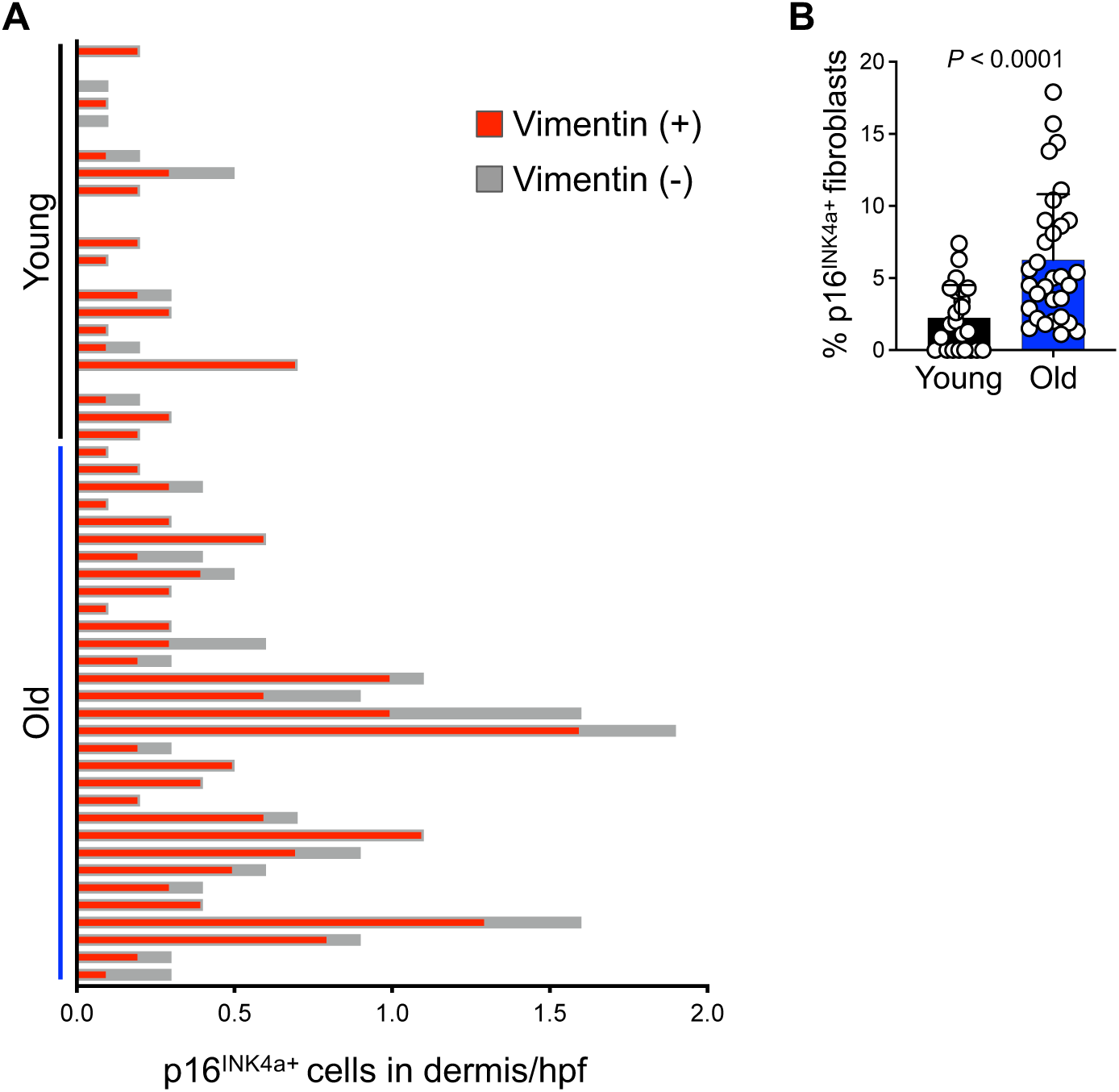
Senescent fibroblasts constitute most of the senescent cells in the dermis of human skin. (**A**) Vimentin positive/negative dermal p16^INK4a+^ cells in each young and old human skin sample. (**B**) Percentage of p16^INK4a^ positive senescent fibroblasts of total fibroblasts per skin sample. Cells are counted blindly and averaged across 10 randomly selected hpf per skin sample. The bar graph shows mean + SD. *n* = 23 in young and *n* = 31 in the old skin group (Mann-Whitney *U* test).

**Supplementary Figure 2.**
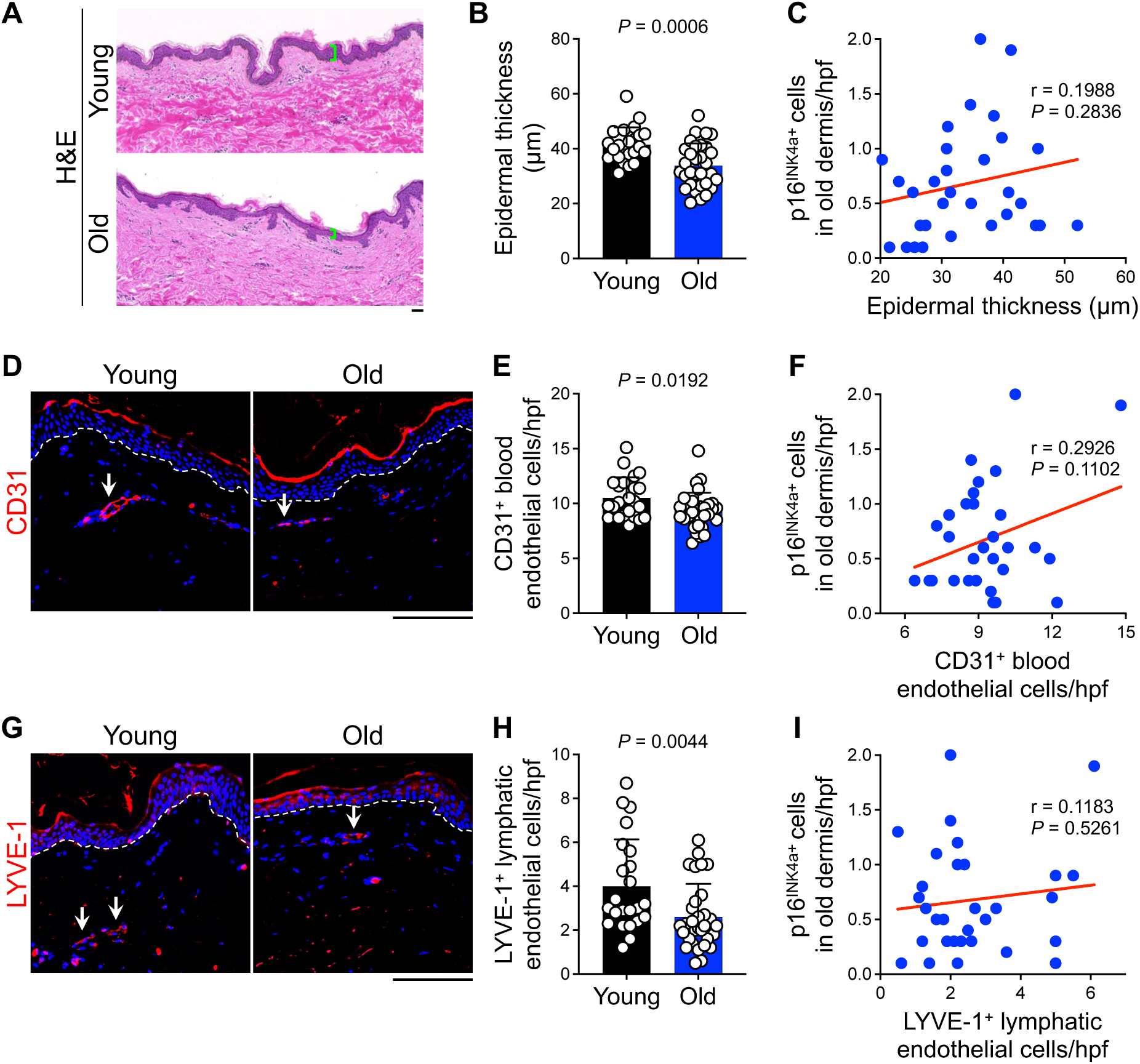
Epidermal thickness and density of blood/lymphatic vessels are reduced with age, but they do not correlate with the number of senescent dermal cells in the old skin. (**A**) Representative hematoxylin and eosin (H&E) staining of young and old skin samples. Epidermal thickness is highlighted by green brackets. (**B**) Quantification of the epidermal thickness (Mann-Whitney *U* test). (**C**) Correlation between the epidermal thickness and the number of dermal p16^INK4a+^ cells in the old skin samples (Student’s *t*-test for the Pearson correlation coefficient). (**D**) Representative IF staining of CD31 (red, a marker for blood endothelial cells) in young and old skin samples. Arrows point to CD31^+^ blood endothelial cells in the dermis. (**E**) The number of dermal CD31^+^ blood endothelial cells per hpf (Mann-Whitney *U* test). (**F**) Correlation between the number of dermal CD31^+^ blood endothelial cells and the number of dermal p16^INK4a+^ cells in the old skin samples (Student’s *t*-test for the Pearson correlation coefficient). (**G**) Representative IF staining of LYVE-1 (red, a marker for lymphatic endothelial cells) in young and old skin samples. Arrows point to LYVE-1^+^ lymphatic endothelial cells in the dermis. (**H**) The number of dermal LYVE-1^+^ lymphatic endothelial cells per hpf (Mann-Whitney *U* test). (**I**) Correlation between the number of dermal LYVE-1^+^ lymphatic endothelial cells and the number of dermal p16^INK4a+^ cells in the old skin samples (Student’s *t*-test for the Pearson correlation coefficient). Nuclei are stained with DAPI (blue), dotted lines in IF images mark the epidermal basement membrane. Cells are counted blindly and averaged across 10 randomly selected hpf per skin sample. Bar graphs show mean + SD. *n* = 23 in young and *n* = 31 in the old skin group. Scale bars: 100 μm.

**Supplementary Figure 3.**
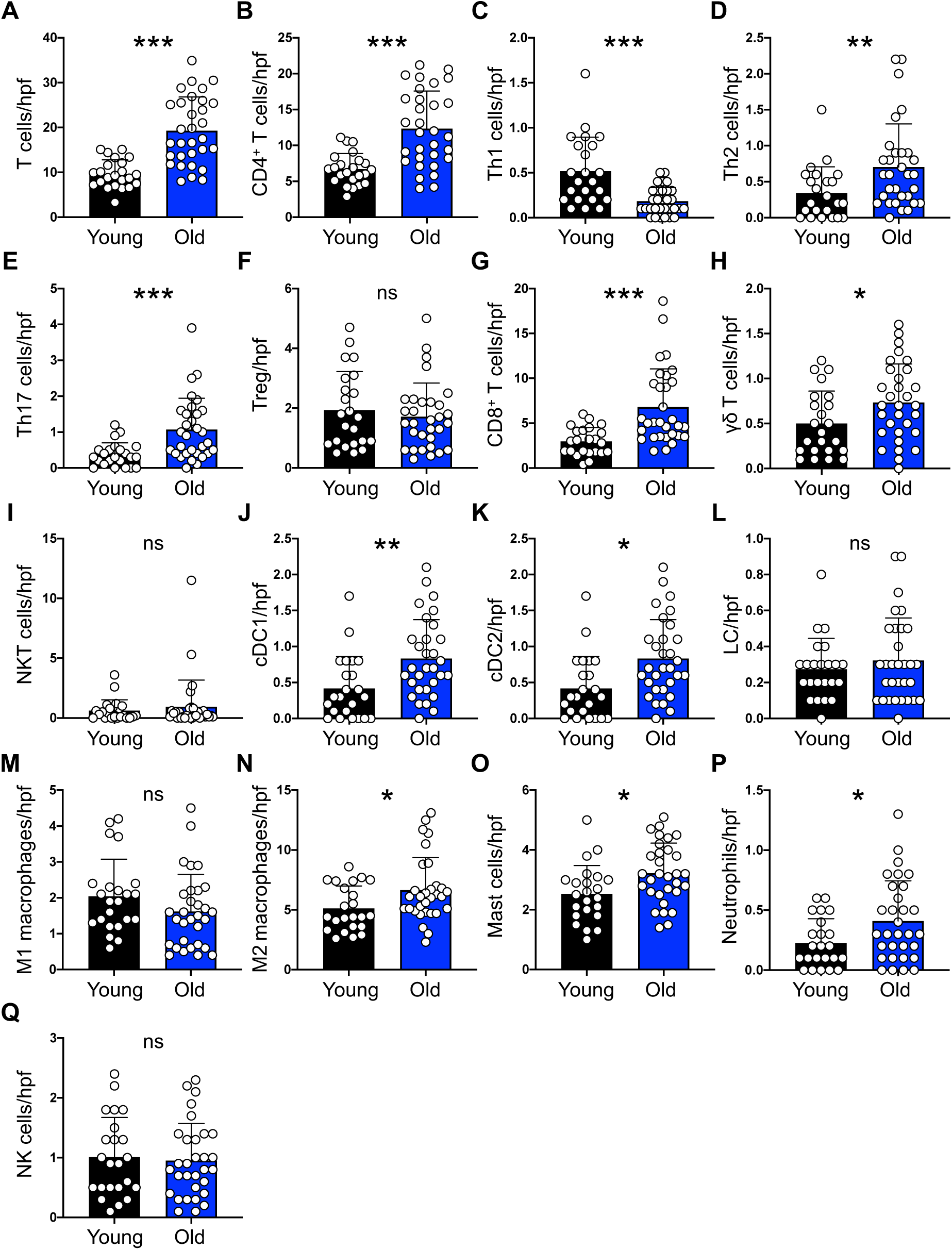
Immune cell quantification in the human skin. (**A**) The number of dermal CD3^+^ cells indicative of T cells per hpf. (**B**) The number of dermal CD3^+^ CD4^+^ cells indicative of CD4^+^ T cells per hpf. (**C**) The number of dermal CD4^+^ T-bet^+^ cells indicative of Th1 cells per hpf. (**D**) The number of dermal CD4^+^ GATA3^+^ cells indicative of Th2 cells per hpf. (**E**) The number of dermal CD4^+^ ROR-γt^+^ cells indicative of Th17 cells per hpf. (**F**) The number of dermal CD4^+^ Foxp3^+^ cells indicative of regulatory T cells (Treg) per hpf. (**G**) The number of dermal CD3^+^ CD8^+^ cells indicative of CD8^+^ T cells per hpf. (**H**) The number of dermal CD3^+^ TCR γδ^+^ cells indicative of γδ T cells per hpf. (**I**) The number of dermal CD3^+^ CD56^+^ cells indicative of NKT cells per hpf. (**J**) The number of dermal CD11c^+^ CD141^+^ cells indicative of cDC1 per hpf. (**K**) The number of dermal CD1c^+^ CD11c^+^ CD207^-^ cells indicative of cDC2 per hpf. (**L**) The number of dermal CD1a^+^ CD207^+^ cells indicative of Langerhans cells (LC) per hpf. (**M**) The number of dermal CD68^+^ CD86^+^ cells indicative of M1 macrophages per hpf. (**N**) The number of dermal CD68^+^ CD206^+^ cells indicative of M2 macrophages per hpf. (**O**) The number of dermal mast cell tryptase^+^ cells indicative of mast cells per hpf. (**P**) The number of dermal neutrophil elastase^+^ cells indicative of neutrophils per hpf. (**Q**) The number of dermal CD3^-^ CD56^+^ cells indicative of NK cells per hpf. Cells are counted blindly and averaged across 10 randomly selected hpf per skin sample. Note that these data are shown as a heatmap in Fig. 2A. Bar graphs show mean + SD. *n* = 23 in young and *n* = 31 in the old skin group. * *p* < 0.05, ** *p* < 0.01, and *** *p* < 0.0001, ns: not significant, Mann-Whitney *U* test.

**Supplementary Figure 4.**
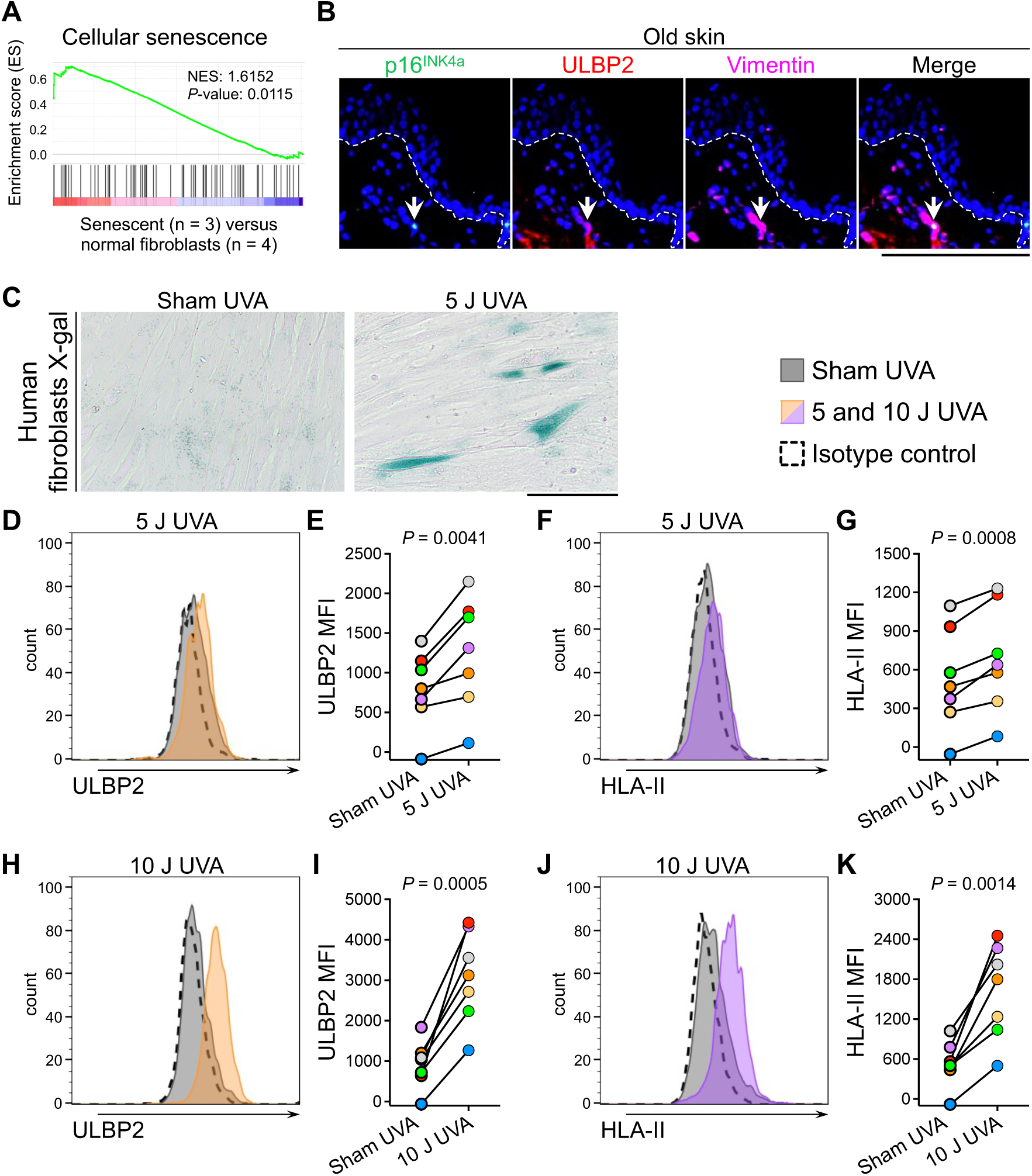
ULBP2 and HLA-II are expressed on the surface of senescent fibroblasts. (**A**) The gene set enrichment analysis plot of cellular senescence in replication-induced senescent dermal fibroblasts (*n* = 3) compared with normal dermal fibroblasts (*n* = 4) from RNA-Seq data. (**B**) Representative IF staining of p16^INK4a^ (green), ULBP2 (red), and vimentin (magenta) in the old skin. The arrows point to a ULBP2^+^ senescent fibroblast in the dermis. Nuclei are stained with DAPI (blue), dotted lines mark the epidermal basement membrane. (**C**) Representative X-gal staining of human dermal fibroblasts irradiated with 5 J of UVA versus sham. (**D**, **E**) Representative flow cytometry histogram (D) and median fluorescence intensity (MFI) (E) of ULBP2 on the surface of dermal fibroblasts one day after irradiation with 5 J of UVA versus sham. (**F**, **G**) Representative flow cytometry histogram (F) and MFI (G) of HLA-II on the surface of dermal fibroblasts one day after irradiation with 5 J of UVA versus sham. (**H**, **I**) Representative flow cytometry histogram (H) and MFI (I) of ULBP2 on the surface of dermal fibroblasts one day after irradiation with 10 J of UVA versus sham. (**J**, **K**) Representative flow cytometry histogram (J) and MFI (K) of HLA-II on the surface of dermal fibroblasts one day after irradiation with 10 J of UVA versus sham. *n* = 7 pairs of adult human dermal fibroblasts, paired *t*-test. Scale bars: 100 μm.

**Supplementary Figure 5.**
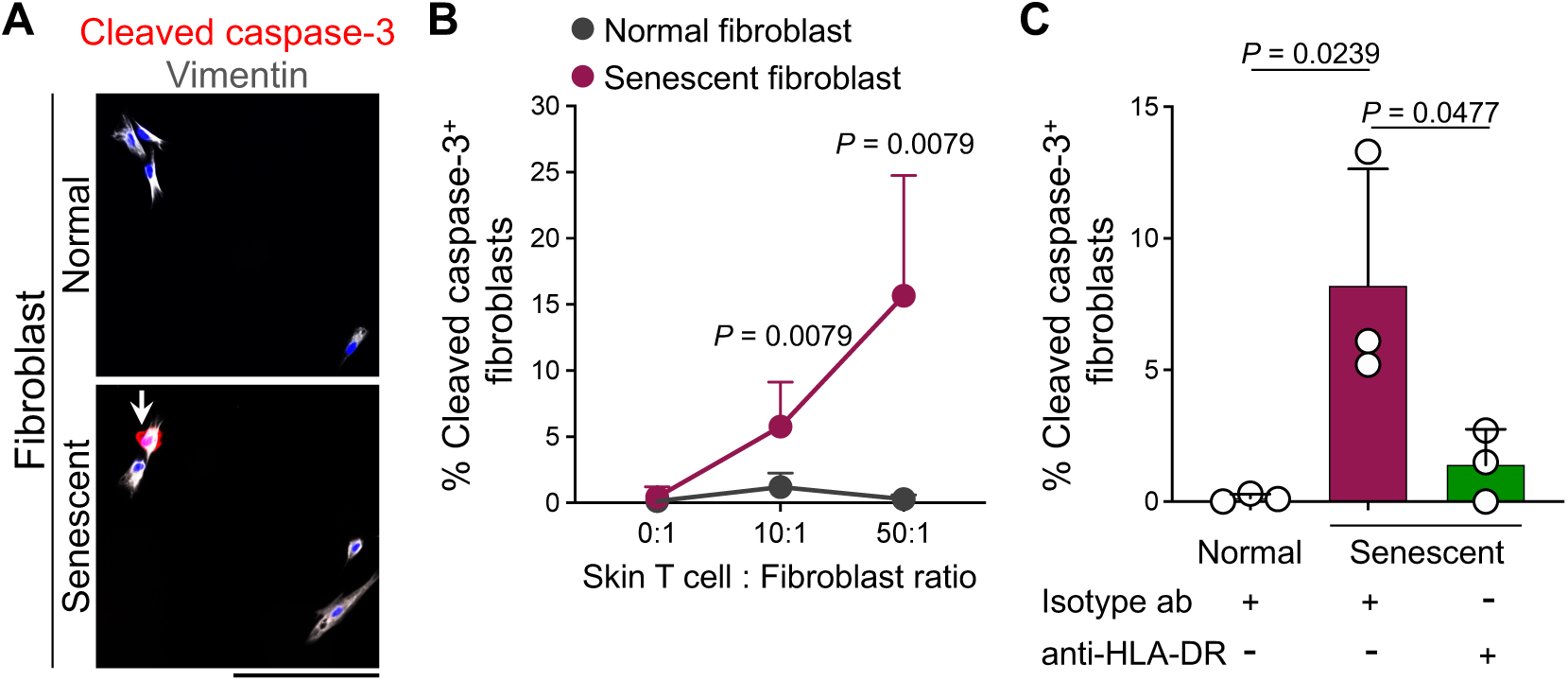
Allogeneic skin-resident T cells selectively eliminate senescent fibroblasts in an HLA-II-dependent manner. (**A**) Representative ICC staining of cleaved caspase-3 (red) and vimentin (gray) in the normal and senescent human dermal fibroblasts after co-culture with skin-derived allogeneic T cells. Nuclei are stained with DAPI (blue). Arrow points to a cleaved caspase-3^+^ apoptotic fibroblast. (**B**) Frequency of cleaved caspase-3^+^ normal and senescent fibroblasts after co-culture with skin-derived allogeneic T cells for 6 h, using a skin T cell to fibroblast ratio of 0:1, 10:1 and 50:1 (*n* = 5 individual T cell donors, Mann-Whitney *U* test). (**C**) Frequency of cleaved caspase-3^+^ normal or senescent fibroblasts after co-culture with skin-derived allogeneic T cells for 6 h. Fibroblasts were pre-incubated with pan-HLA-DR blocking or isotype control antibody for 18 h. Skin-derived T cell to fibroblast ratio of 50:1 (*n* = 3 individual T cell donors, one-way ANOVA). Cells are counted blindly and averaged across 10 randomly selected hpf per sample. Bar graphs show mean + SD. Scale bar: 100 μm.

**Supplementary Figure 6.**
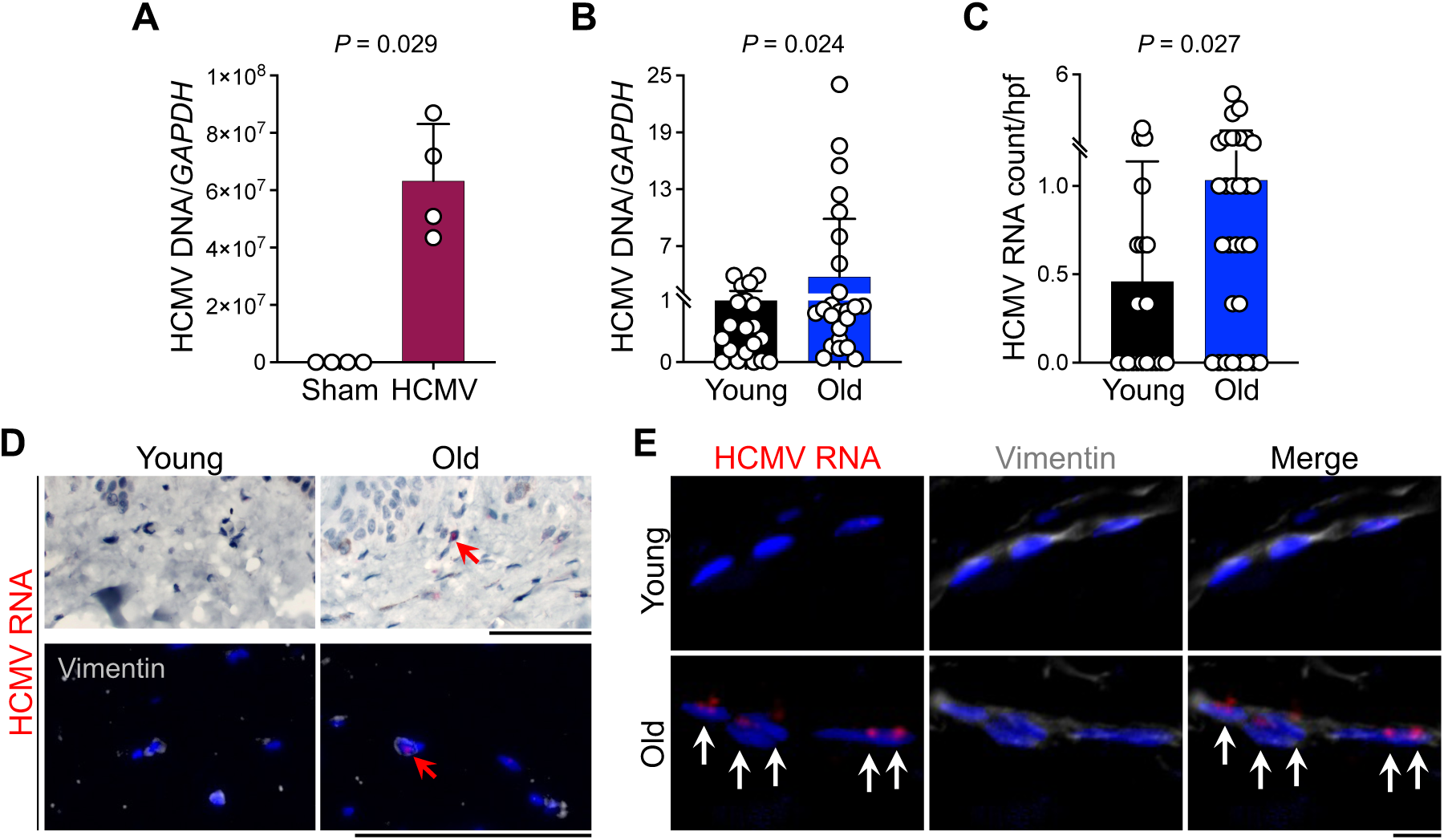
HCMV DNA and RNA expression are upregulated in the old human skin. (**A**) Quantitative PCR for HCMV DNA detection in HCMV-versus sham-infected human fetal fibroblasts using HCMV *UL83* primers. Data are presented as the ratio of *GAPDH* expression (*n* = 4 per group). (**B**) Quantitative PCR for HCMV DNA detection in the young versus old human skin samples using HCMV *UL83* primers. Data are presented as the ratio of *GAPDH* expression. (**C**) Quantification of HCMV RNA *in situ* hybridization (ISH) count in the young versus old skin samples. HCMV RNA ISH positive signals are counted blindly and averaged across three hpf with HCMV RNA ISH signal if any per skin sample. (**D**) Representative images of HCMV RNA ISH (red, arrow) immunohistochemistry staining and IF staining with vimentin (gray) in the dermis of young and old skin (scale bar: 100 μm). (**E**) Representative images of HCMV RNA ISH (red) with IF staining of vimentin (gray) of dermal fibroblasts in the young and old skin. Arrows point to HCMV RNA^+^ signals in fibroblasts (scale bar: 10 μm). HCMV RNA ISH probe is designed to detect *UL123* (IE1) transcript. Nuclei are stained with DAPI (blue). Bar graphs show mean + SD. *n* = 21 in young and *n* = 30 in the old skin group. Mann-Whitney *U* test.

**Supplementary Figure 7.**
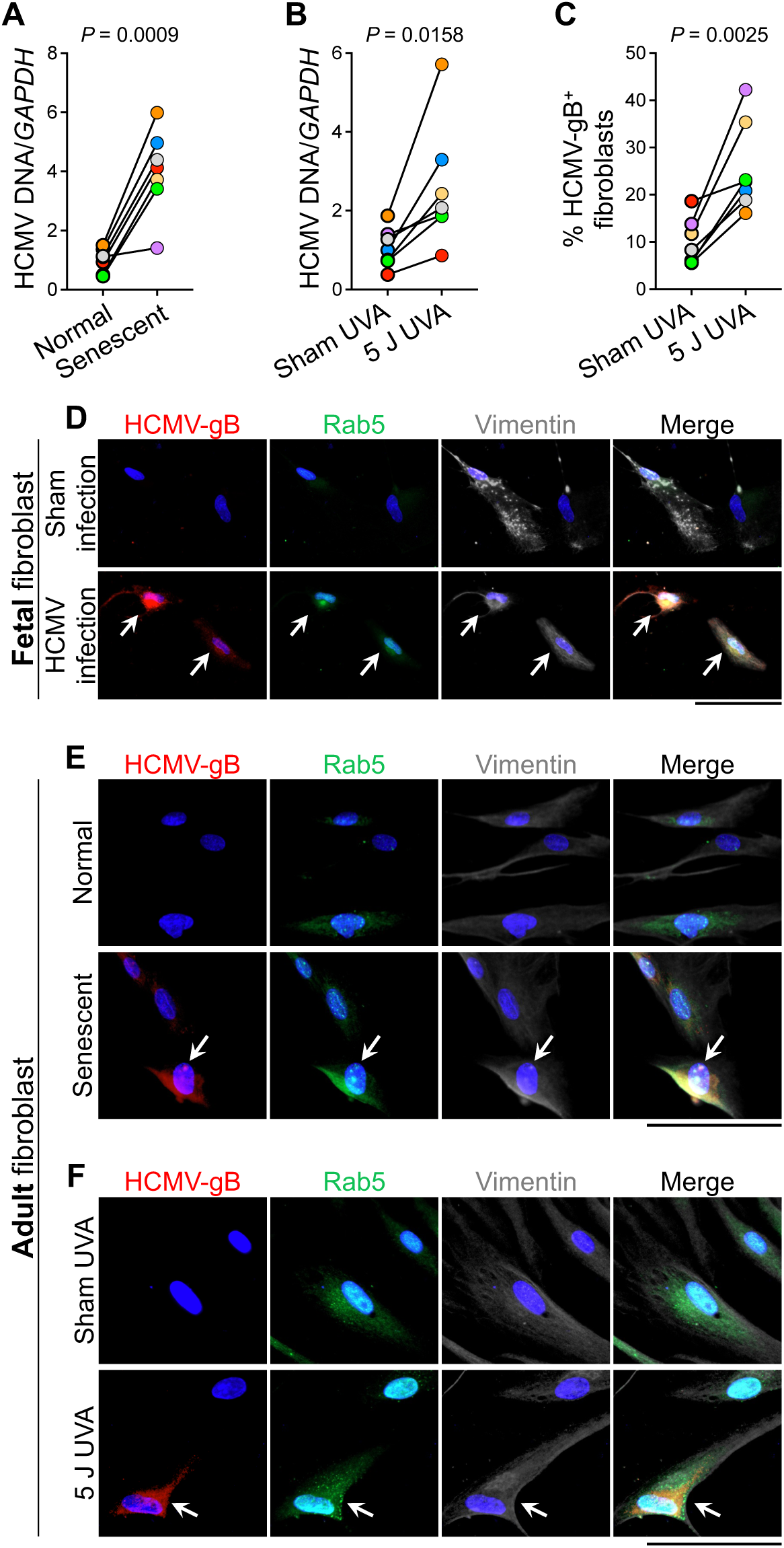
HCMV-gB expression is induced by HCMV infection of fetal fibroblasts, and repeated replication and UVA radiation of adult fibroblasts. (**A**) Quantitative PCR for HCMV DNA detection in normal and replication-induced senescent fibroblasts using HCMV *UL83* primer. Data are presented as the ratio of *GAPDH* expression. (**B**) Quantitative PCR for HCMV DNA detection in fibroblasts one day after 5 J UVA versus sham UVA radiation using HCMV *UL83* primer. Data are presented as the ratio of *GAPDH* expression. (**C**) Frequency of HCMV-gB^+^ fibroblasts five days after 5 J UVA versus sham UVA radiation. (**D**) Representative ICC staining of HCMV-gB (red), Rab5 (green), and vimentin (gray) in HCMV-versus sham-infected human fetal fibroblasts. Arrows point to HCMV-gB and Rab5 signal co-localization in fibroblasts. (**E**) Representative ICC staining of HCMV-gB (red), Rab5 (green), and vimentin (gray) in normal and replication-induced senescent fibroblasts. Arrows point to HCMV-gB and Rab5 signal co-localization in a fibroblast. (**F**) Representative ICC staining of HCMV-gB (red), Rab5 (green), and vimentin (gray) in fibroblasts five days after 5 J UVA or sham UVA radiation. Arrows point to HCMV-gB and Rab5 signal co-localization in a fibroblast. Nuclei are stained with DAPI (blue). *n* = 7 pairs of adult human dermal fibroblasts, paired *t*-test. Scale bars: 100 μm.

**Supplementary Figure 8.**
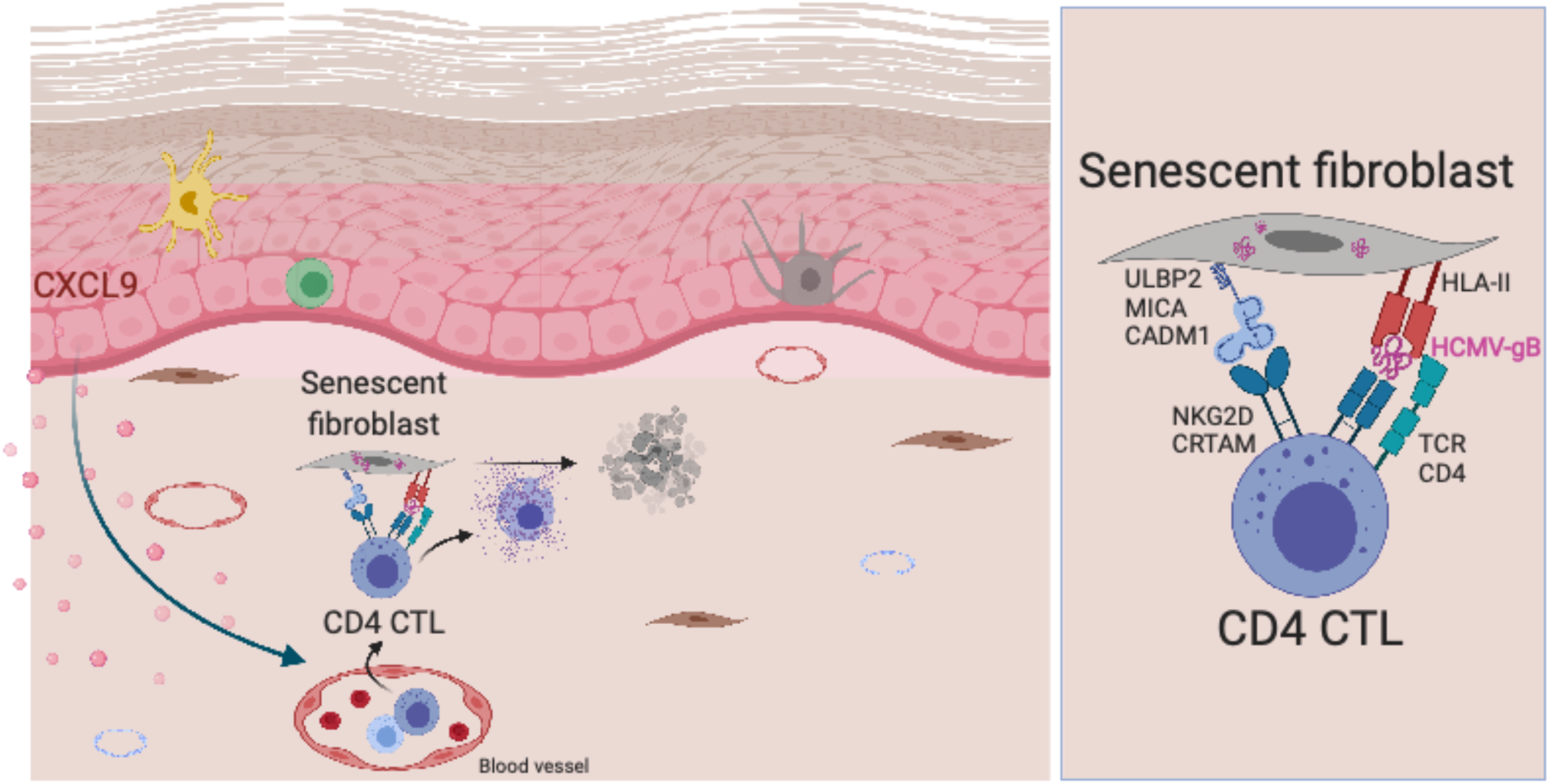
Schematic diagram of HCMV-gB-mediated clearance of senescent fibroblasts by cytotoxic CD4 CTL. Created with BioRender.com

### SUPPLEMENTARY TABLES

**Supplementary Table 1.** Human skin samples information.

|  | Young skin<br>(n=23) | Old skin<br>(n=31) | *P value* |
| --- | --- | --- | --- |
| Age, years |  |  |  |
| Mean (SD) | 23.1 (3.4) | 62.1 (5.9) | < 0.0001 |
| Range | 16-28 | 53-74 |  |
| Gender, number (%) |  |  | - |
| Male | 0 (0) | 0 (0) |  |
| Female | 23 (100) | 31 (100) |  |
| Epidermal thickness, mean (SD), $\mu\text{m}$ | 41.5 (6.3) | 33.8 (2.7) | 0.0006 |
| No. of hair follicle units per mm skin, mean (SD) | 0.19 (0.16) | 0.15 (0.16) | 0.346 |

**Supplementary Table 2.**
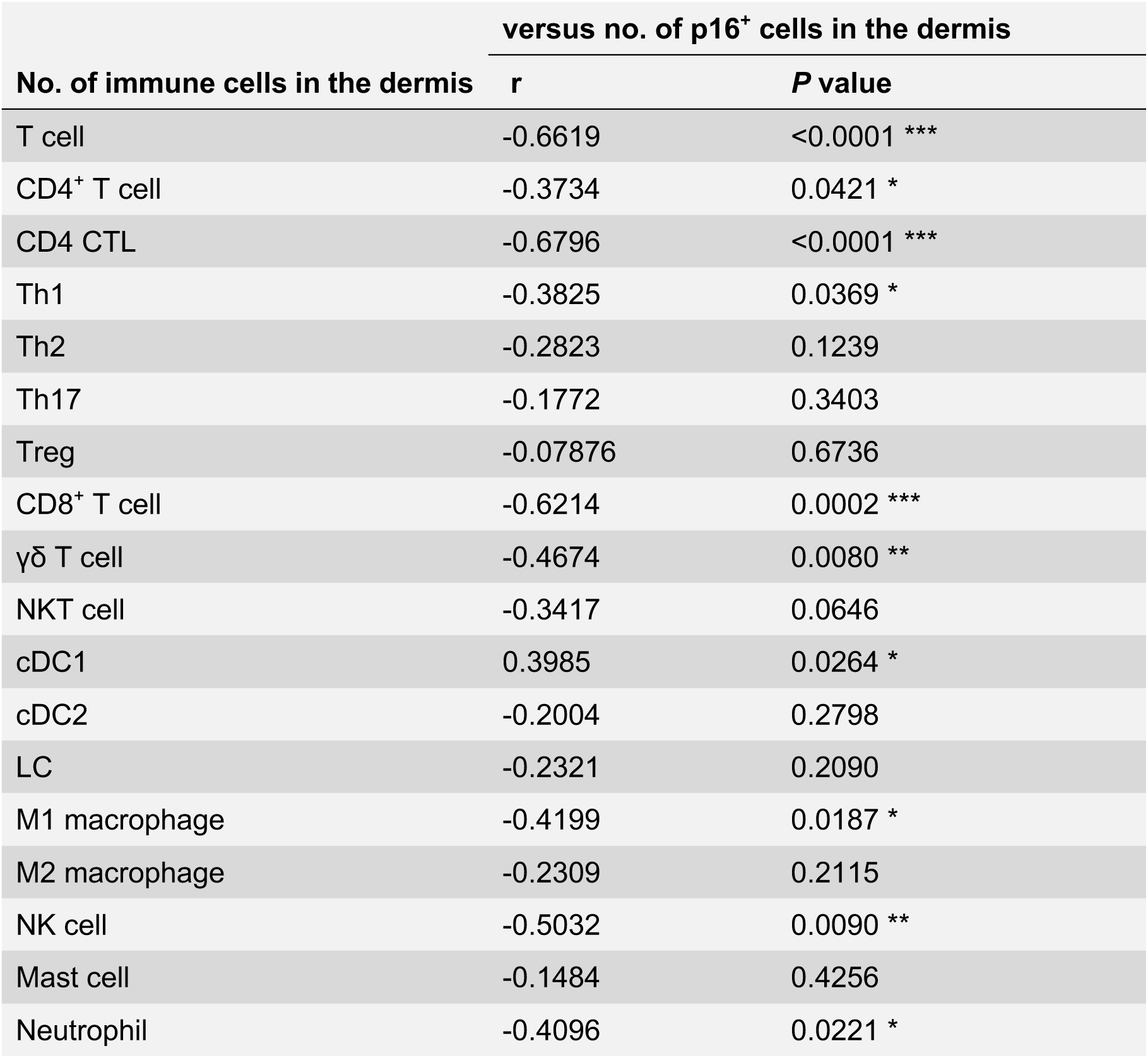
Immune cell correlation with senescent (p16^+^) cell accumulation in the dermis of the old human skin samples.

| No. of immune cells in the dermis | versus no. of p16<sup>+</sup> cells in the dermis |  |
| --- | --- | --- |
|  | r | P value |
| T cell | -0.6619 | <0.0001 *** |
| CD4<sup>+</sup> T cell | -0.3734 | 0.0421 * |
| CD4 CTL | -0.6796 | <0.0001 *** |
| Th1 | -0.3825 | 0.0369 * |
| Th2 | -0.2823 | 0.1239 |
| Th17 | -0.1772 | 0.3403 |
| Treg | -0.07876 | 0.6736 |
| CD8<sup>+</sup> T cell | -0.6214 | 0.0002 *** |
| γδ T cell | -0.4674 | 0.0080 ** |
| NKT cell | -0.3417 | 0.0646 |
| cDC1 | 0.3985 | 0.0264 * |
| cDC2 | -0.2004 | 0.2798 |
| LC | -0.2321 | 0.2090 |
| M1 macrophage | -0.4199 | 0.0187 * |
| M2 macrophage | -0.2309 | 0.2115 |
| NK cell | -0.5032 | 0.0090 ** |
| Mast cell | -0.1484 | 0.4256 |
| Neutrophil | -0.4096 | 0.0221 * |

**Supplementary Table 3.**
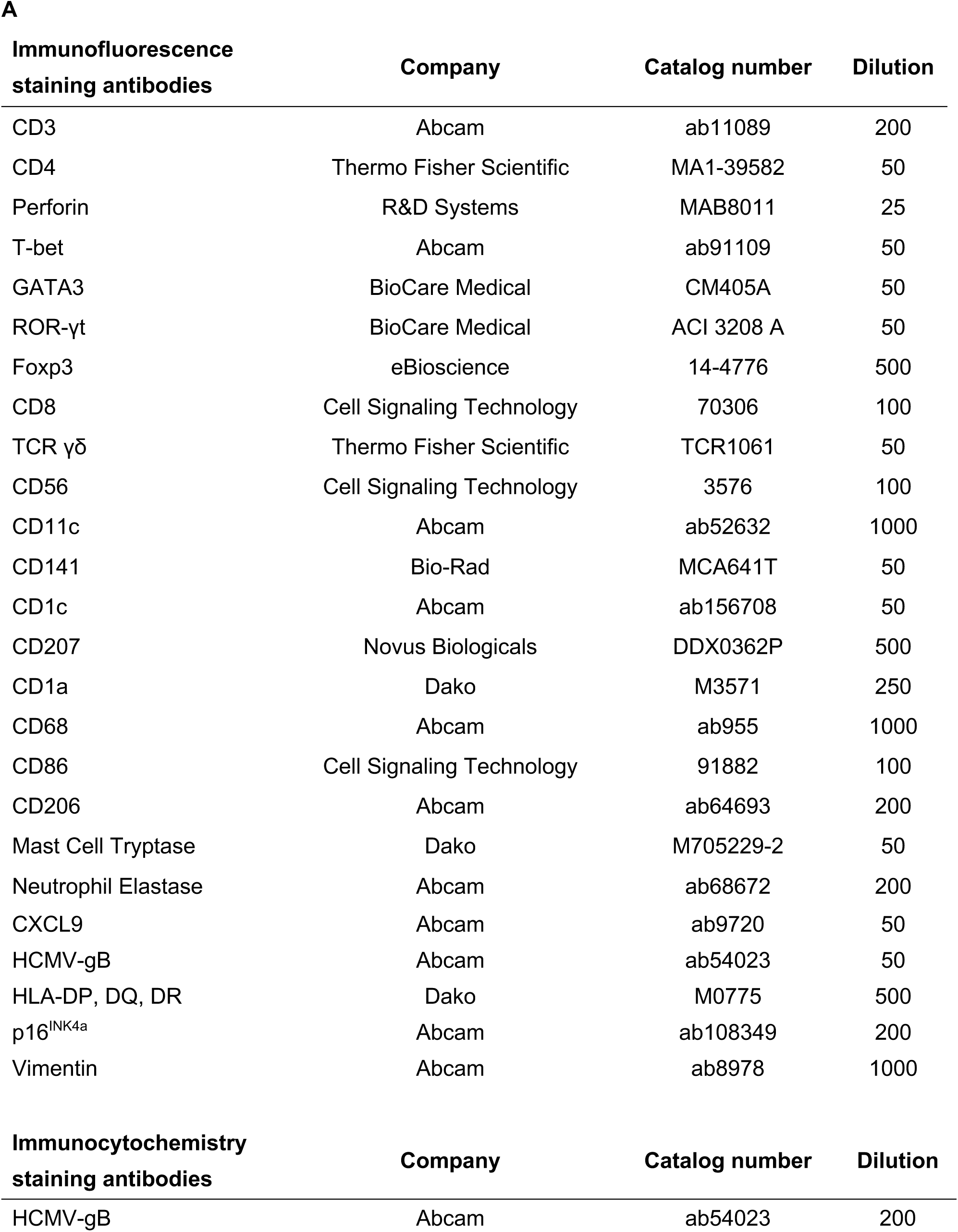
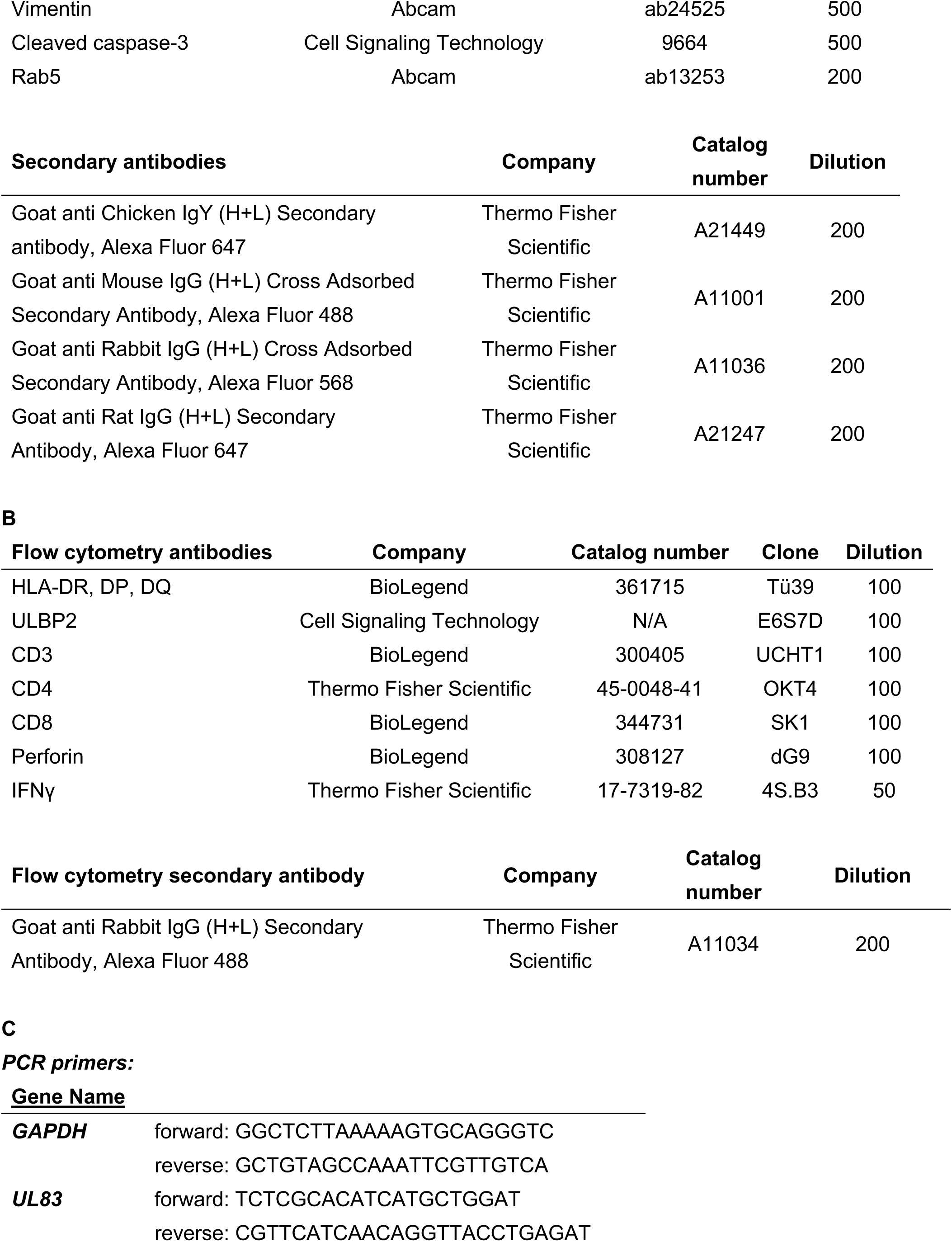
Antibodies for immunofluorescence and immunocytochemistry (A), antibodies for flow cytometry (B), and primers for qPCR (C).

